# An orthogonal near-infrared optical switch for wireless neuromodulation in freely behaving mice

**DOI:** 10.64898/2026.04.12.717984

**Authors:** Zheyu Xie, Limin Pan, Yipeng Hua, Bojun Hou, Peishan Xiang, Yi Wu, Shifang Shan, Ximei Yan, Yuhan Chen, Peizhe Gao, Jiulin Du, Jianan Liu

## Abstract

Modulating neuronal activity with light is a powerful tool for neuroscience research. However, currently available technologies often require invasive fibers for delivery of visible light, causing tissue damage and limiting behavioral studies. Although near-infrared (NIR) neuronal modulation improves tissue penetration depth, sustained NIR illumination during neuromodulation raises significant concerns about photothermal effects. Here, we introduce a dual-wavelength near-infrared switch (Dual-NIR Switch) that uses two transcranial NIR inputs to initiate and terminate neuronal activation, enabling tether-free neuromodulation in freely moving mice. Dual-NIR Switch employs orthogonal dichromatic upconversion nanoparticles that emit blue and green light under 980 nm and 808 nm excitation, respectively, to activate and inactivate the step-function opsin SOUL. Therefore, transient 980 nm NIR illumination initiates neuronal excitation, which will remain excited without further stimulation but will be rapidly terminated on demand by a subsequent 808 nm illumination. Upon transcranial 980 nm and 808 nm NIR illuminations in freely moving mice, we achieve on-demand control of behavioral paradigms across tunable timescales, ranging from seconds to minutes and even extending to sub-hour durations. By eliminating the need for sustained NIR irradiation, Dual-NIR Switch offers an on-demand, duration-tunable neuromodulation tool for both basic neuroscience and potential therapeutic applications in treating brain diseases.

## Introduction

Optical neuromodulation offers unparalleled advantages for interrogating neural circuits, combining precise cell-type specificity with high spatiotemporal resolution^1–3^. A central challenge in deploying this powerful methodology, however, lies in the minimally invasive delivery of light to deep brain regions for behaviorally relevant studies. Conventional approaches rely on implanted optical fibers or head-mounted hardware, which can cause local tissue damage and provoke chronic neuroinflammatory responses^4^. Moreover, tethering and head-mounted hardware can restrict long-term mobility and alter naturalistic behaviors, introducing confounds that complicate behavioral interpretation^5^. Therefore, these limitations motivate the development of less invasive, wireless strategies for delivering light into brain.

Significant efforts have focused on the development of microscale light-emitting diodes, which can supply visible light directly to targeted brain regions^6–9^, yet their chronic utility is often constrained by issues of mechanical reliability, power delivery, and heat dissipation. To enable transcranial optical neuromodulation without intracranial surgery, opsin engineering has shifted their excitation spectra toward longer wavelength^10,11^. Recently, the deep tissue penetration of near-infrared (NIR) light has been leveraged for transcranial neuromodulation through NIR-triggered nanotransducers^12–16^. Such nanotransducers include upconversion nanoparticles^17–20^ that convert transcranial NIR into visible emission to drive opsins, and photothermal nanoagents^21,22^ that translate NIR light into localized heat to activate thermosensitive channels. Despite these advances, a fundamental constraint persists: neuromodulation is maintained only under continuous external NIR irradiation, thereby coupling the duration of neuronal activity directly to the period of NIR irradiation. Consequently, the operating window of such NIR-based technologies is typically restricted to seconds, as prolonged NIR illumination raises cumulative thermal risks. Therefore, there is a pressing need for a NIR-based neuromodulation platform that minimizes thermal burden to enable duration-tunable behavioral control.

Here, we introduce a dual-wavelength NIR switch (Dual-NIR Switch) that decouples the state of neuronal activity from continuous NIR irradiation. Dual-NIR Switch pairs orthogonal dichromatic upconversion nanoparticles (odUCNPs) with the SOUL, a step-function opsin^23^ opened by blue light and closed by green light. Upon NIR illumination at 980 nm, blue emission from odUCNPs activates SOUL to initiate neuronal activation, which persists for up to ∼30 min without further irradiation. This persistent activity can be terminated on demand by a subsequent 808 nm illumination, which elicits green emission of odUCNPs to rapidly inactivate SOUL. In freely moving mice, Dual-NIR Switch enables fully wireless control of behaviors with durations ranging from seconds to sub-hour timescale. Owing to its excellent biocompatibility and negligible thermal load, Dual-NIR Switch offers a transformative approach for chronic, wireless neuroscience studies.

## Design and development of Dual-NIR Switch

We engineered odUCNPs with a multilayer core-shell structure to achieve orthogonal emission of blue and green light under distinct NIR excitations^24–27^ (Figure 1a). In our design, the luminescent core was co-doped with Yb³⁺, Er³⁺, and a low concentration of Nd³⁺ (0.5 mol%) to minimize concentration quenching^28^. The first shell layer, doped with 20 mol% Nd³⁺ as a sensitizer, was then incorporated to efficiently harvest 808 nm photons for green upconversion emission. Additionally, Yb³⁺ and Tm³⁺ were co-doped into a shell layer to enable blue emission under 980 nm excitation. To prevent undesired energy transfer and surface-related quenching effects, inert NaYF₄ shells are grown between the two luminescent centers and on the outermost layer, respectively (Figure 1b). Sequential shell growth yielded monodisperse, hexagonal phase odUCNPs with an average diameter of ∼40 nm (Figure 1e and Figure S1). Elemental mapping confirmed the spatially segregated incorporation of lanthanide dopants (Yb³⁺/Tm³⁺ and Nd³⁺/Er³⁺) into separate shell layers, which allows for orthogonal emission (Figure 1f and Figure S2).

**Figure 1.**
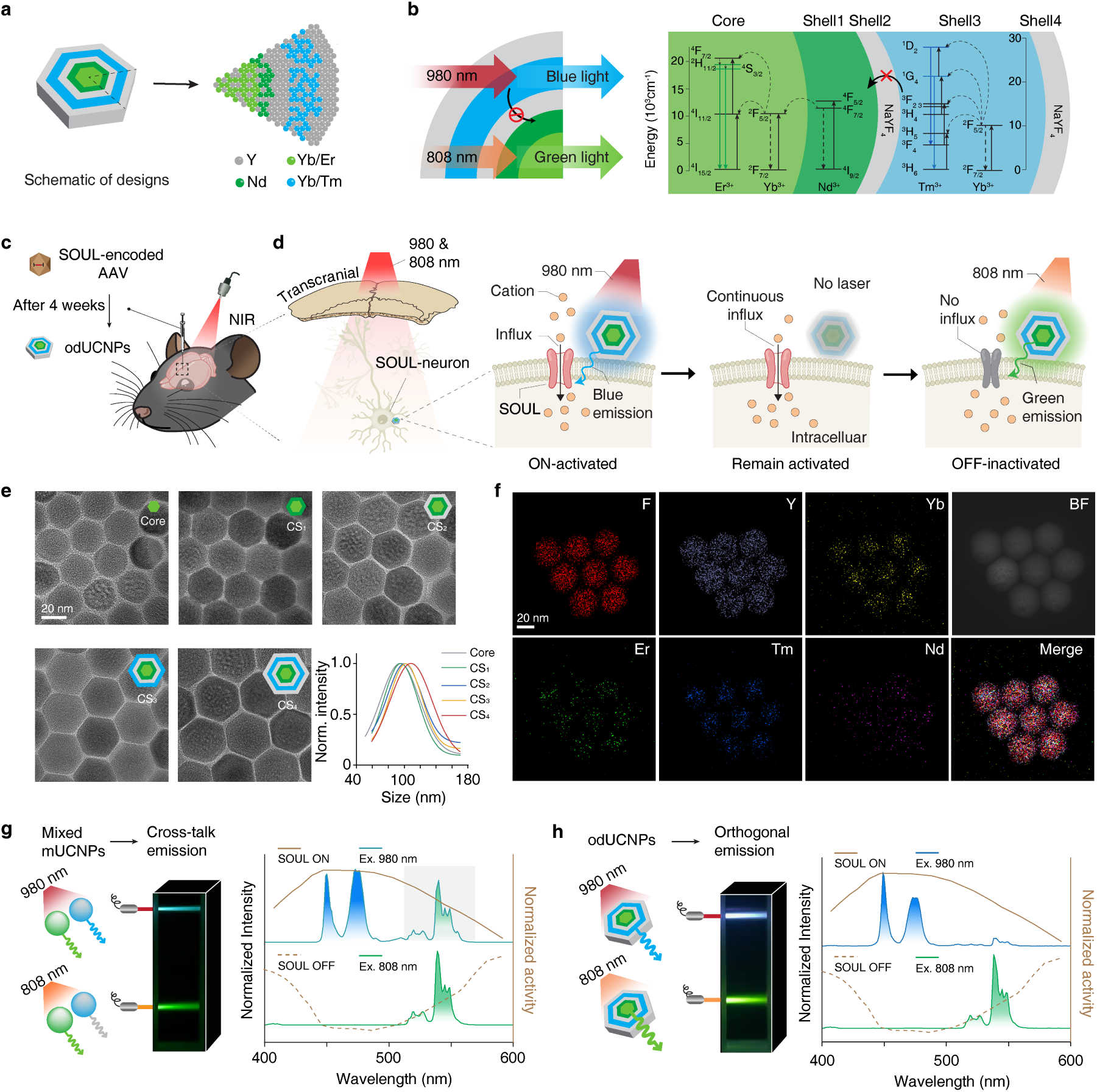
Development of Dual-NIR Switch. (a) Core-multishell structure of odUCNPs with spatially segregated lanthanide dopants (for example, Yb³⁺/Er³⁺ in the core and Yb³⁺/Tm³⁺ in an outer shell) to enable orthogonal emission. (b) Orthogonal emission scheme under distinct NIR excitations: 980-nm excitation yields predominantly blue emission (Tm³⁺-centered), whereas 808-nm excitation yields predominantly green emission (Er³⁺-centered), with minimal spectral crosstalk. (c) In vivo workflow: delivery of SOUL-encoding AAV followed by local administration of odUCNPs in the targeted brain region. (d) Working principle of Dual-NIR Switch: 980-nm-driven blue upconversion activates SOUL (ON state), producing sustained neuronal excitation (≥30 min), whereas 808-nm-driven green upconversion deactivates SOUL (OFF state) and rapidly terminates excitation. (e) TEM images acquired after successive shell-growth steps and corresponding normalized size distributions. Images share a same scale bar. (f) EDS elemental maps confirming the designed composition and dopant segregation. Images share a same scale bar. (g) and (h) Comparison of mixed mUCNPs (g) and odUCNPs (h): photographs and emission spectra under 980-nm and 808-nm excitation overlaid with SOUL activation (top) and deactivation (bottom) action spectra; grey shading highlights undesired crosstalk emission from mUCNPs under 980-nm excitation, which is suppressed in odUCNPs.

While monochromatic UCNPs (mUCNPs) can efficiently activate neuronal activity in combination with opsins, a physical mixture of two types of mUCNPs with distinct emissions fails to achieve orthogonal dichromatic emission due to inevitable interparticle crosstalk (Figure 1g). Consequently, such mixtures are incapable of selectively tuning the opening and closing of SOUL. In contrast, the odUCNPs developed here integrate orthogonal dichromatic emission within a single nanoparticle, which can emit blue light (¹G4 → ³H₆, ¹D₂ → ³F₄ transitions) under 980 nm excitation and green light (²H₁₁/₂ → ⁴I₁₅/₂, ⁴S₃/₂ → ⁴I₁₅/₂ transitions) under 808 nm excitation, respectively (Figure 1b). This light output precisely matches the activation and inactivation spectra of the step-function opsin SOUL^29^ (Figure 1h).

Leveraging orthogonal emission of odUCNPs, Dual-NIR Switch enables remote, transcranial neuromodulation in freely moving mice, without wires or head-mounted hardware and, crucially, without requiring sustained NIR illumination. This Dual-NIR Switch was achieved by sequential microinjections of SOUL-encoding adeno-associated virus (AAV) and odUCNPs into the target brain region (Figure 1c). A brief 980 nm illumination prompts the odUCNPs to emit blue light, activating the step-function opsin SOUL and thereby inducing a depolarizing cation influx. Once activated, SOUL remains in an active state without any light illumination, maintaining neuronal excitation for up to ∼30 min. Whenever needed, a short 808 nm illumination elicits green emission, inactivating SOUL and precisely terminating neuronal activity (Figure 1d).

## Dual-NIR Switch modulates cell activities in vitro

To validate whether Dual-NIR Switch can orthogonally control SOUL activation and inactivation, we performed whole-cell patch-clamp recordings on 293T cells expressing SOUL (Figure 2a–c). The odUCNPs were rendered hydrophilic and membrane-affinitive via glutathione modification^30,31^ (Figure S3), promoting their adhesion to the cell surface (Figure 2d and Figure S4), thereby ensuring efficient light delivery to membrane-bound SOUL.

**Figure 2.**
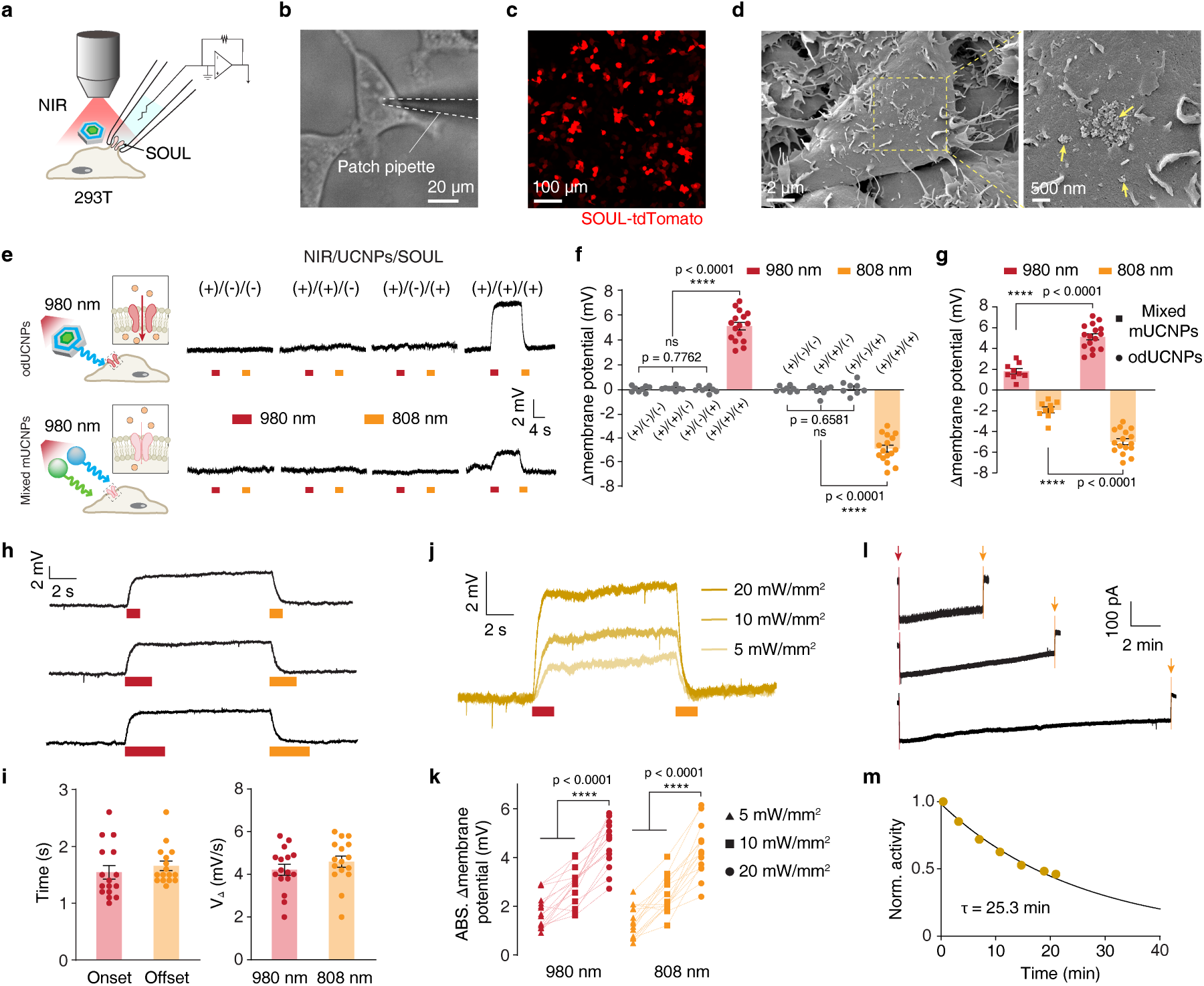
Dual-NIR Switch modulates cell activities *in vitro*. (a) Patch-clamp configuration for SOUL-expressing 293T cells incubated with odUCNPs and stimulated upon sequential NIR illuminations. (b) Brightfield image of 293T cells during whole-cell patch-clamp recording. (c) Confocal image of SOUL–tdTomato expression of 293T cells. (d) SEM images of 293T cells after incubation showing odUCNPs attached to the plasma membrane (arrows); right, magnification view. (e) Representative voltage traces from cells incubated with mixed mUCNPs or odUCNPs upon sequential NIR illumination at 980 nm (red) followed by 808 nm (orange). (f) Quantification of NIR-illumination-induced membrane potential changes under indicated conditions. Data are shown as mean ± s.e.m. (n = 16 cells for (+)/(+)/(+) and n = 8 cells for others, one-way ANOVA with Tukey’s multiple comparison test). (g) Quantification of NIR-illumination-induced membrane potential changes between mUCNPs and odUCNPs. Data are shown as mean ± s.e.m. (n = 9 cells for mixed mUCNPs and n = 16 cells for odUCNPs, t-test). (h) Representative voltage traces showing responses to varying durations of NIR illumination (1, 3, 5 s). (i) Onset/offset delay (left) and rate of NIR-illumination-induced membrane potential changes (right). Data are shown as mean ± s.e.m. (n = 16 cells). (j) Representative voltage traces at indicated NIR power densities. (k) NIR-illumination-induced absolute membrane potential changes as a function of NIR power density. (n = 15 cells, one-way ANOVA with Tukey’s multiple comparison test). (l) Representative current traces showing sustained activation after 980 nm illumination (red arrow) and deactivation by 808 nm illumination (orange arrows). (m) Normalized current decay after cessation of 980-nm stimulation fitted with a single-exponential function (τ = 25.3 min).

Systematic testing across four experimental conditions confirmed significant membrane potential shifts only in the presence of SOUL, odUCNPs, and NIR illumination (Figure 2e). Under this condition, 980 nm illumination induced a depolarization of 5.10 ± 0.30 mV (mean ± s.e.m.), and subsequent 808 nm illumination caused a repolarization of 4.99 ± 0.29 mV. Such responses markedly exceed changes all other control groups (Figure 2f).

To highlight the importance of the orthogonal emission of odUCNPs in opening and closing the SOUL, we compared odUCNPs against a physical mixture of mUCNPs: one (NaYF₄:Yb/Tm) emitting blue light upon 980 nm illumination and the other (NaYF₄:Yb/Nd/Er) emitting green light upon 808 nm illumination (Figure S5a,b). While either mUCNPs alone could activate or inactivate SOUL (Figure S5c–f), however, their mixture exhibited significantly reduced response amplitude and slower kinetics compared to odUCNPs (Figure 2g and Figure S5g–i). This impairment stems from inter-particle spectral crosstalk in the mixture, which introduces competing emissions and weakens precise neuromodulation. In contrast, the designed odUCNPs effectively avoid this limitation (Figure 1g,h).

Dual-NIR Switch also exhibited well-defined temporal characteristics, with activation and inactivation time constants of 1.54 ± 0.12 s and 1.66 ± 0.08 s at 20 mW/mm² (Figure 2h,i). The membrane potential response followed a clear dose-dependent relationship with NIR power (Figure 2j,k), enabling duration tunable modulation. Critically, Dual-NIR Switch supported long term modulation without sustained energy input. SOUL activation persisted for minutes following a brief 980 nm illumination, with a current decay time constant of 25.3 min, consistent with its intrinsic biophysics (Figure 2l,m and Figure S6). Activity was reliably and fully terminated on demand by a subsequent 808 nm illumination.

## Dual-NIR Switch wirelessly modulates short-duration behavior

To examine whether this Dual-NIR Switch can timely modulate mouse behavior via activating neurons in vivo, considering its capabilities for orthogonally modulating membrane potential, an AAV encoding SOUL was injected into the secondary motor cortex (M2), followed four weeks later by microinjection of odUCNPs at the same site (Figure 3a). Robust SOUL expression was confirmed (Figure 3b), and bio-TEM revealed odUCNPs localized near cell bodies and axons (Figure 3c), ensuring efficient light delivery to membrane-bound SOUL. Notably, a detectable number of odUCNPs persisted in the brain, indicating their good tissue retention and potential for long-term functionality after a single administration (Figure S7).

**Figure 3.**
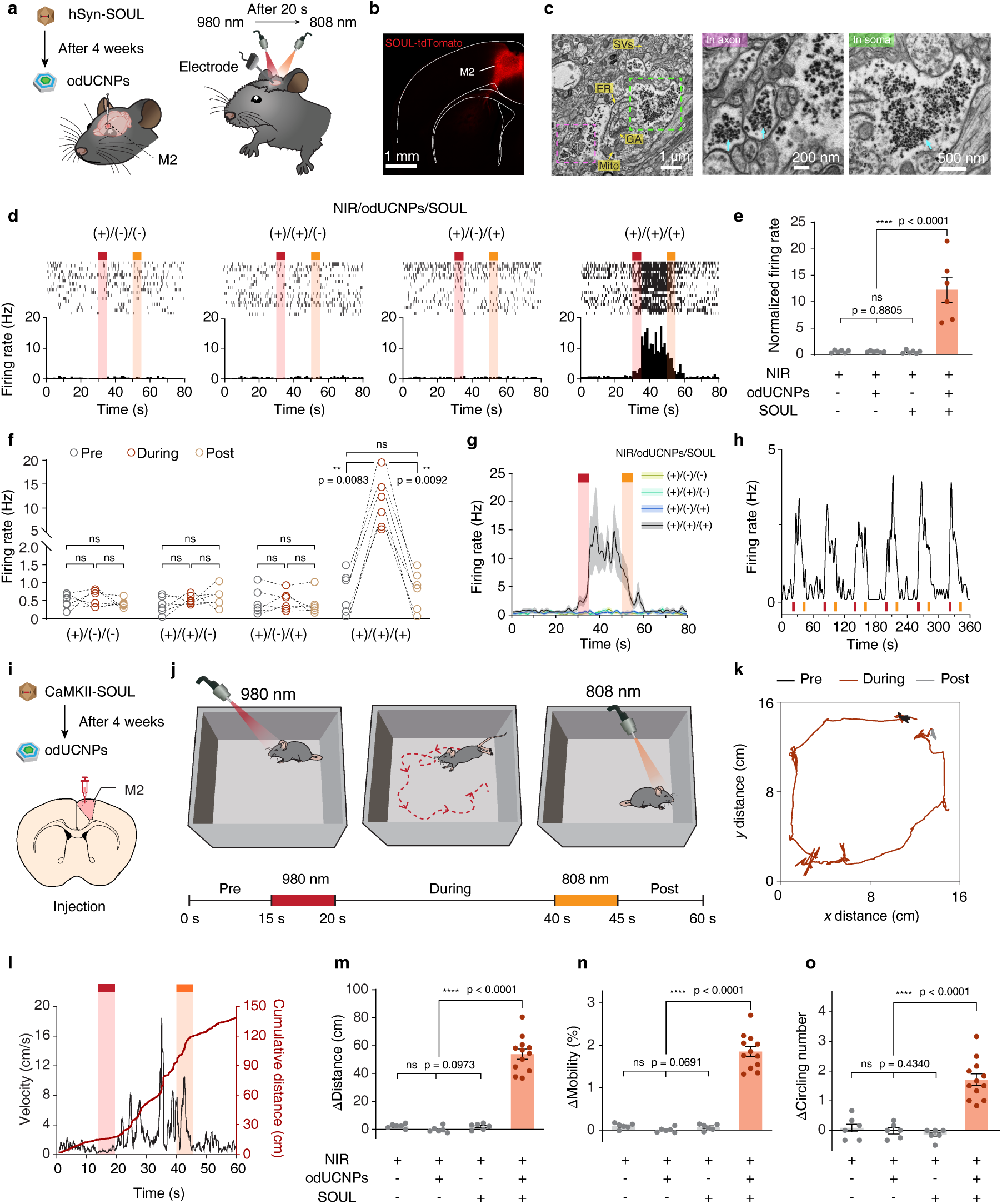
Modulation of short-duration locomotion behavior in M2 brain region by Dual-NIR Switch. (a) Schematic of the injection timeline and in vivo electrophysiological recordings of M2 under modulation by Dual-NIR Switch. (b) Confocal image showing SOUL expression in M2. (c) Bio-TEM images showing the distributions of odUCNPs in M2. Magnified views highlight clusters in axon and soma. (d) Raster plots and corresponding firing rate in response to sequential NIR illumination at 980 nm followed by 808 nm under indicated conditions. Each row of the raster plot represents a single trial. (n = 6 mice, N = 18 trails). (e) Quantification of normalized firing rate within “During” period under indicated conditions. Data are shown as mean ± s.e.m. (f) Quantification of the firing rate before 980 nm illumination (Pre, 0–15 s), after 808 nm illumination (Post, 65–80 s) and in between (During, 35–50 s). In (e) and (f), each point/point-set indicates a unit from one mouse averaged over 3 trials, n = 6 mice for each group, one-way ANOVA with Tukey’s multiple comparison test. Full set p-values are provided in Supporting Table. (g) Average firing rate dynamics under indicated conditions. Data are shown as mean ± s.e.m. (h) Representative neuronal firing rate dynamics following cycles of neuromodulation by Dual-NIR Switch. (i) and (j) Experimental timeline (i) and NIR stimulation pattern (j) for the locomotion enhancement test. (k) Representative trajectory of a mouse before 980 nm illumination (Pre, 0–15 s), in between (During, 15–45 s) and after 808 nm illumination (Post, 45–60 s). (l) Kinetics of instant velocity (black) and cumulative distance (red) from a representative trail. (m)–(o) Changes in distance moved (m), mobility (n) and circling number (o) between “During” period and baseline (sum or average of “Pre” and “Post” period), respectively. Data are shown as mean ± s.e.m. (n = 12 mice for (+)/(+)/(+), n = 6 mice for others; one-way ANOVA with Tukey’s multiple comparison test).

In vivo multichannel electrophysiology recording in anesthetized mice showed that neuronal firing rates during the interval between the two NIR illuminations exhibited a positive correlation with increasing power density (Figure S8a–d). Between the 980 nm and 808 nm illuminations at 24 mW/mm², neuronal firing rates significantly increased exclusively in NIR+/odUCNPs+/SOUL+ group (Figure 3d–f). Notably, the Dual-NIR switch enabled neuromodulation with an onset latency of 0.5 s and an offset latency of 1.5 s, and is repeatable over multiple cycles of 980 nm and 808 nm illumination (Figure 3g,h and Figure S8e,f), demonstrating a timely tunable and reliable approach for neuronal activation.

We then tested whether this neuronal activation in M2 could translate to wireless behavioral modulation in freely moving mice^32,33^. Following unilateral SOUL expression and odUCNPs delivery in M2 (Figure 3i), mice received sequential NIR illumination at 980 nm followed by 808 nm (5 s, 40 mW/mm²) in an open field (Figure 3j). Locomotion was reliably enhanced by Dual-NIR Switch between two NIR illuminations (Video S1), with increased velocity, distance moved, and mobility specifically (Figure 3k–n and Figure S9, S10a). Unilateral activation of M2 neurons also induced circling behaviour^34,35^, as evidenced by NIR+/odUCNPs+/SOUL+ group showing significantly more turns and higher angular velocity than any other controls (Figure 3o and Figure S10b).

Together, these results demonstrate that Dual-NIR Switch enables wireless, switchable control of neuronal activity and short-duration locomotor behavior in freely moving mice, decoupling sustained neuronal excitation from continuous external energy input.

## Dual-NIR Switch wirelessly modulates medium-duration behavior

We next examined whether the Dual-NIR Switch could achieve transcranial neuromodulation over minutes in deep brain regions. Thus, we targeted glutamatergic neurons in the lateral hypothalamic area (LHA), a key feeding center whose activation suppresses food intake via downstream inhibition of dopaminergic signalling^36,37^. Following bilateral expression of SOUL in LHA neurons, odUCNPs were injected at the same region four weeks later (Figure 4a and Figure S11).

**Figure 4.**
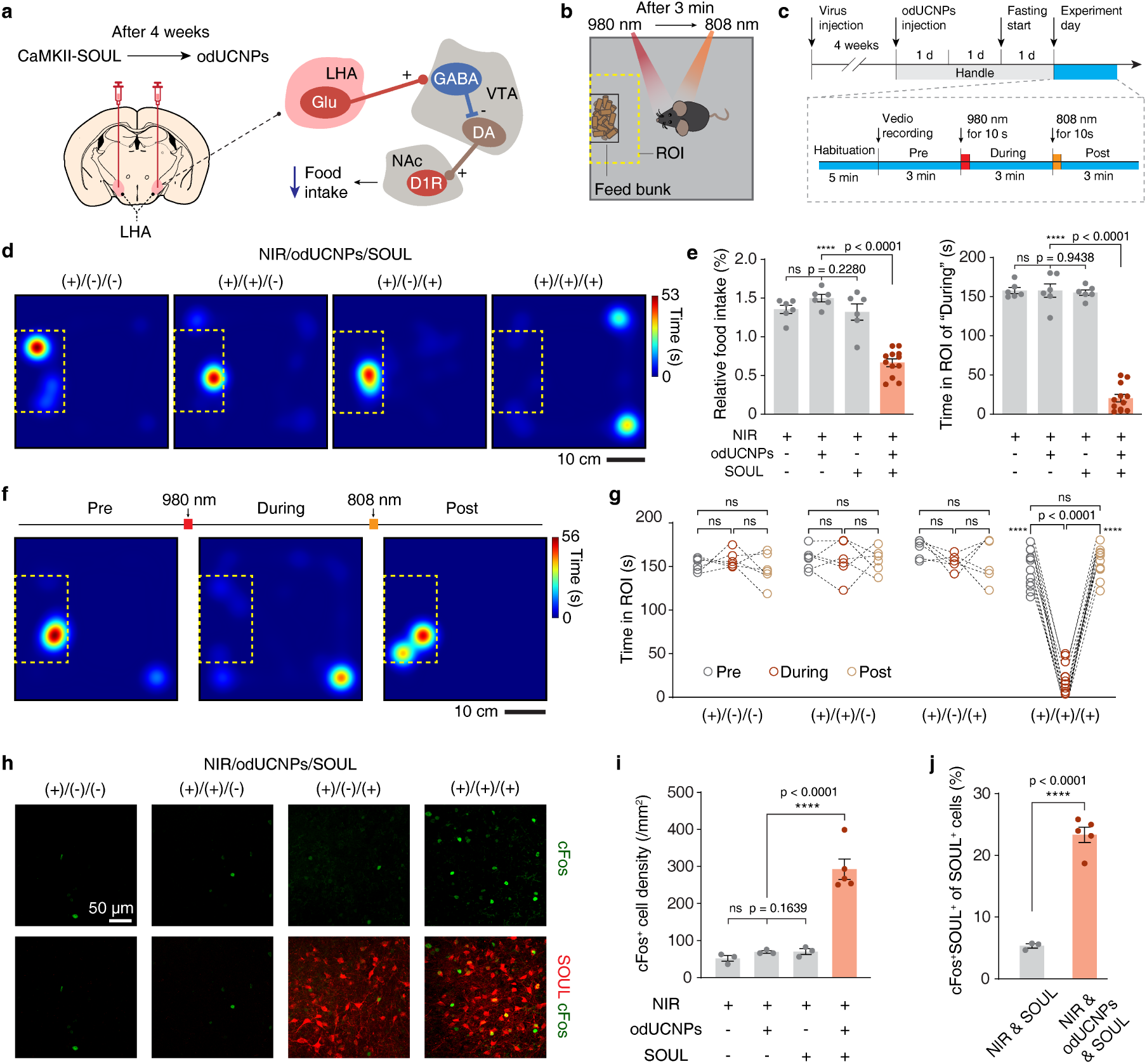
Modulation of medium-duration feeding behavior in a deep-brain region LHA by Dual-NIR Switch. (a) Experimental scheme and circuit context for free-access feeding: SOUL and odUCNPs were targeted to LHA glutamatergic neurons, whose activation reduces food intake. (b) Arena schematic: the feeding zone (yellow dashed box) was defined as the region of interest (ROI). (c) Behavioral timeline: after habituation, mice were subjected to sequential NIR illumination at 980 nm followed by 808 nm (10 s each at 50 mW/mm², CW). (d) Representative heatmaps of “During” period under indicated conditions. (e) Quantification of relative food intake (food intake normalized to body weight; left) and time in ROI of “During” period (right), Data are shown as mean ± s.e.m. (f) Representative heatmaps of “Pre”, “During” and “Post” period of one mouse in (+)/(+)/(+) group. (g) Quantification of the time spent in ROI of “Pre”, “During” and “Post” period. Each point-set represents one mouse. In (e) and (g), n = 12 mice for (+)/(+)/(+) and n = 6 mice for others, one-way ANOVA with Tukey’s multiple comparison test. Full set p-values are provided in Supporting Table. (h) c-Fos immunostaining in LHA to validate neuronal activation; SOUL (red), c-Fos (green) and merge. Images share one scale bar. (i) Quantification of density of c-Fos⁺ cells under indicated conditions (one-way ANOVA with Tukey’s multiple comparison test). (j) Specificity of activation: percentage of SOUL-expressing neurons that were also c-Fos⁺ (unpaired t-test). In (i) and (j), data are shown as mean ± s.e.m. Each point represents the average of fluorescence measurements from 3 brain slices, n = 5 mice in (+)/(+)/(+) group and n = 3 for others.

In a real-time feeding test^38^ (Figure 4b,c), all mice received sequential NIR illumination at 980 nm followed by 808 nm (50 mW/mm², CW). Mice in NIR+/odUCNPs+/SOUL+ group were reliably suppressed feeding by Dual-NIR Switch between two NIR illuminations (Video S2). After 980 nm NIR stimulation, these mice promptly left the food zone, leading to a significant reduction in both food intake and feeding duration compared to all control groups (Figure 4d,e and Figure S12). In the meantime, locomotor activity was unaffected, confirming that the observed decrease of food intake was specific to feeding (Figure S13). Feeding resumed to baseline levels after the 808 nm NIR illumination, demonstrating the neuronal activation could be efficiently terminated (Figure 4f,g).

Immunostaining for the immediate-early gene c-Fos showed elevated neuronal activation selectively in the LHA of NIR+/odUCNPs+/SOUL+ mice (Figure 4h,i and Figure S14). The proportion of SOUL-expressing neurons that were c-Fos⁺ increased from 5.3 ± 0.3% (without odUCNPs) to 23.3 ± 1.2% (with odUCNPs), confirming efficient and targeted recruitment of the circuit (Figure 4j). These results demonstrate that Dual-NIR Switch can induce continuous neuromodulation over several minutes in a deep brain region, enabling precise control of motivated behavior.

## Dual-NIR Switch wirelessly modulates long-duration behavior

Building on its ability to modulate behavior over seconds and minutes, we next asked whether the Dual-NIR Switch could support longer-duration, reward-related neuromodulation in deep brain circuits. To this end, we targeted dopaminergic neurons in the ventral tegmental area (VTA), a region that underlies associative learning and typically requires sustained activity to link contexts with reward^39^. SOUL was unilaterally expressed in the VTA, followed by odUCNPs injection four weeks later (Figure 5a).

**Figure 5.**
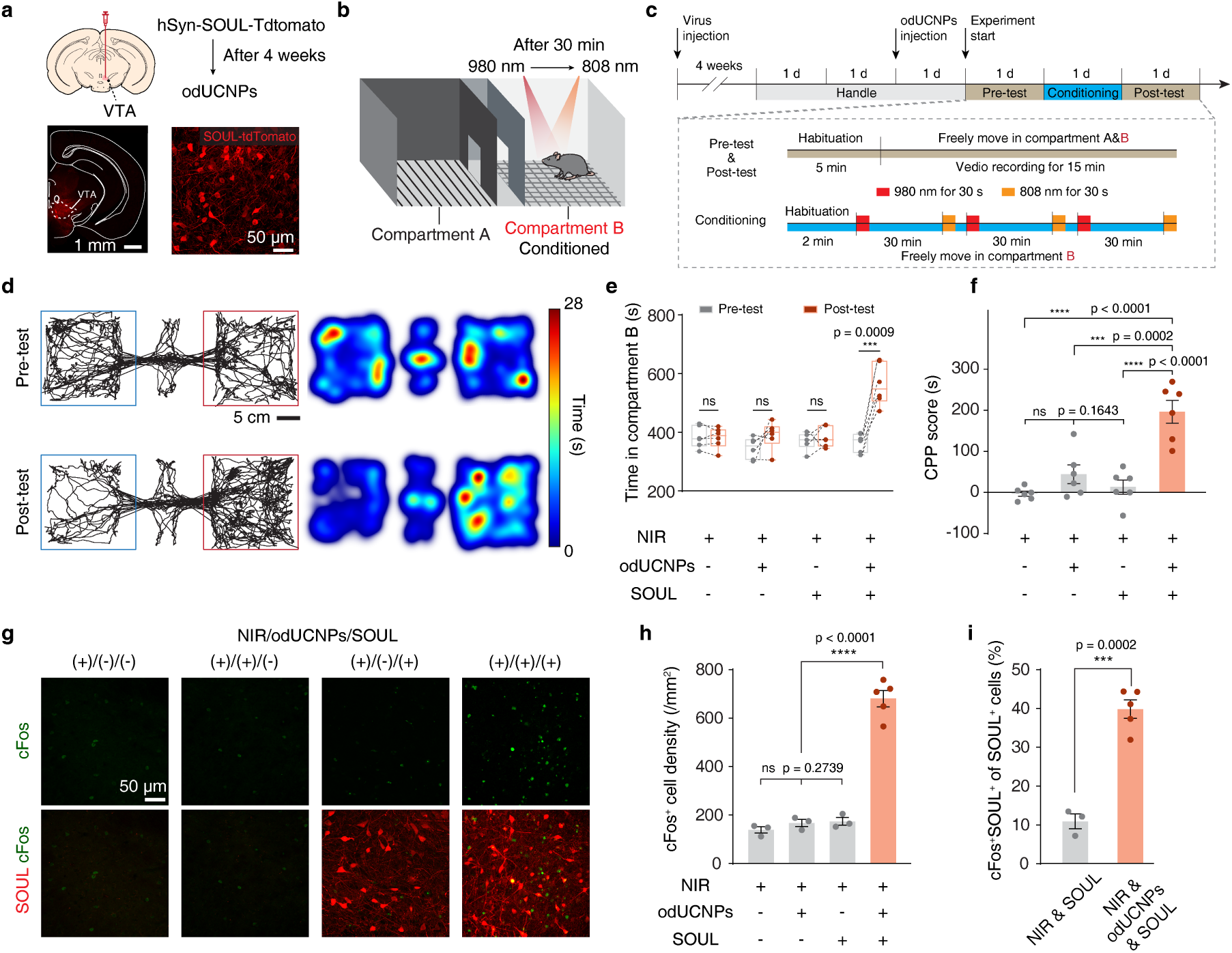
Modulation of long-duration reward conditioning behavior in a deep-brain region VTA by Dual-NIR Switch. (a) Experimental scheme (top) and confocal validation of SOUL expression in VTA (bottom). (b) Schematic of CPP apparatus: compartment B is conditioned by sequential NIR illumination at 980 nm followed by 808 nm. Mid-compartment separates compartments A and B and can be gated. (c) Timeline of CPP test: in the Pre- and Post-test, the mice freely moved in all compartments. In conditioning, mice were confined to compartment B and received sequential NIR illumination. Both 980 nm and 808 nm illumination were applied for 30 seconds at a power density of 60 mW/mm², with a pulse duration of 0.1 second and a repetition rate of 5 Hz. (d) Representative trajectories (left) and the corresponding heatmaps (right) for a mouse in the (+)/(+)/(+) group during the Pre-test and Post-test. (e) Quantification of time spent in compartment B of Pre- and Post-test under indicated conditions (box plots: min–max with interquartile range; n = 6 mice per group; paired t-test; full set p-values are provided in Supporting Table). (f) CPP scores under indicated conditions. Data are shown as mean ± s.e.m. (n = 6 mice per group, one-way ANOVA with Tukey’s multiple comparison test). (g) c-Fos immunostainings in VTA validate neuronal activation; SOUL (red), c-Fos (green) and merge. Images share one scale bar. (h) Quantification of density of c-Fos⁺ cells under indicated conditions (one-way ANOVA with Tukey’s multiple comparison test). (i) Specificity of activation: percentage of SOUL-expressing neurons that were also c-Fos⁺ (unpaired t-test). In (h) and (i), data are shown as mean ± s.e.m. Each point represents the average of fluorescence measurements from 3 brain slices, n = 5 mice in (+)/(+)/(+) group and n = 3 for others.

We employed a conditioned place preference (CPP) paradigm^40,41^ (Figure 5b), which is widely used to evaluate motivational effects of a certain context. After a baseline preference test (Day 1), mice were confined to conditioned compartment (B) on Day 2 and received three sessions of Dual-NIR Switch stimulation (Figure 5c). In the post-test (Day 3), only NIR+/odUCNPs+/SOUL+ mice showed a significant shift towards the conditioned compartment (Figure 5d and Figure S15). Time spent in compartment B increased from 365.9 ± 12.9 s to 562.4 ± 21.9 s, yielding a markedly higher CPP score compared to all controls (Figure 5e,f and Figure S16). In this behavioral test, general locomotor parameters were also unaffected during the stimulation of Dual-NIR Switch (Figure S17).

Immunostaining for the c-Fos confirmed that neuronal activation in the VTA was specific to the complete experimental condition (Figure 5g,h and Figure S18). Furthermore, the proportion of SOUL-expressing neurons that were c-Fos⁺ increased from 10.9 ± 1.9% (without odUCNPs) to 39.8 ± 2.4% (with odUCNPs), demonstrating efficient modulation of targeted circuit (Figure 5i).

These results show that the Dual-NIR Switch can induce behaviorally relevant neuromodulation over sub-hour duration. Unlike conventional NIR-based methods that require continuous irradiation, our technology provides temporally precise control of neuronal activation with minimal cumulative light exposure, offering a flexible methodology for studying long-duration neural processes.

## Biocompatibility of Dual-NIR Switch

The Dual-NIR Switch requires only microinjections of odUCNPs and brief, low-power NIR illumination. We evaluated the biocompatibility of each component by administering them separately and in combination, examining both their individual and combined effects (Figure 6a).

**Figure 6.**
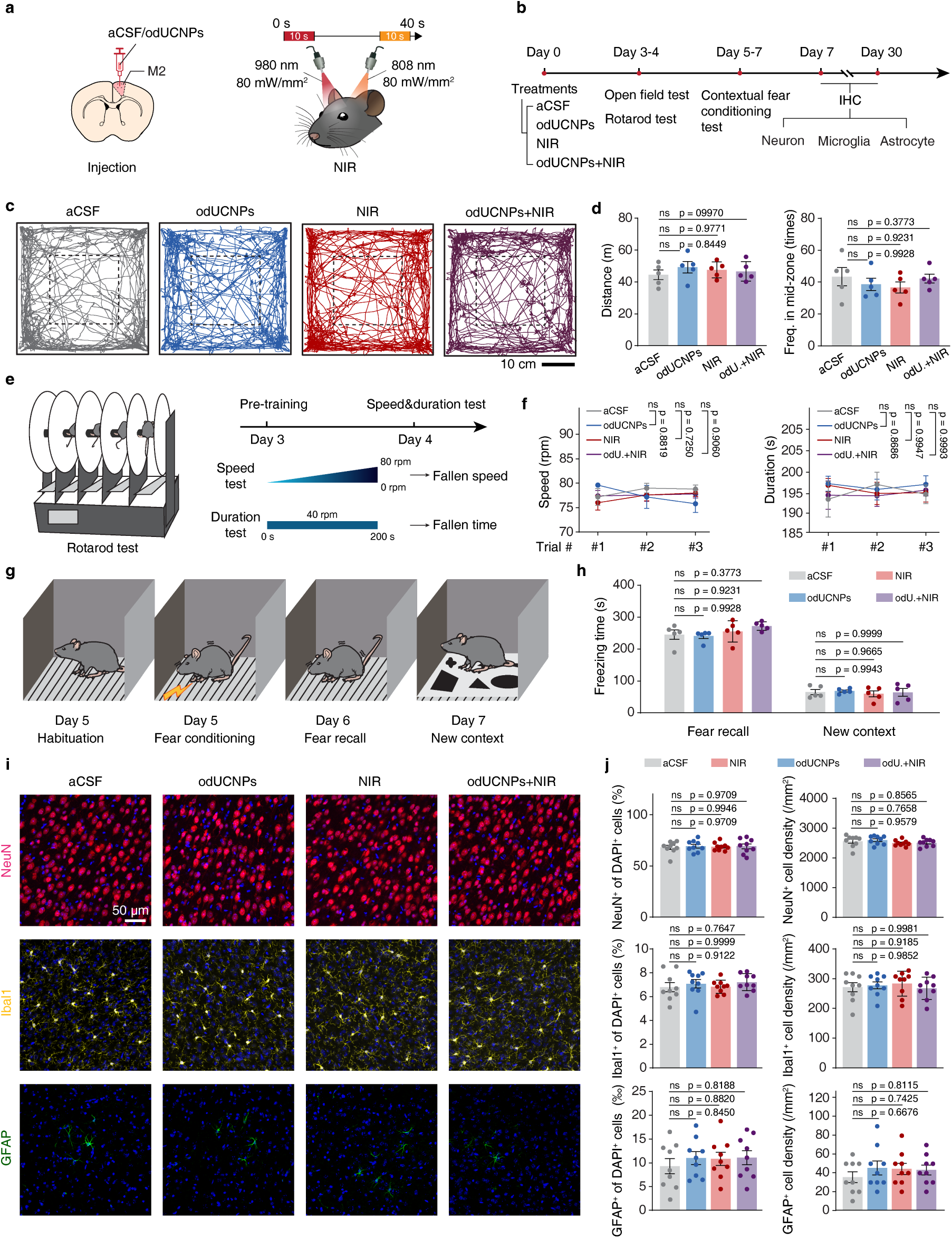
Biocompatibility assessments of Dual-NIR Switch. (a) Treatment paradigms: injection of odUCNPs and NIR illumination patterns. (b) Timeline for biocompatibility assessments: treatments followed by behavioral tests and IHC. (c) Representative open-field trajectories (central zone outlined). (d) Quantification of distance moved (left) and mid-zone entries (right). Data are shown as mean ± s.e.m. (n = 5 mice in each group, one-way ANOVA with Tukey’s multiple comparison test). (e) Rotarod test schematic and timeline. (f) Quantification of rotarod performance (fallen speed and fallen latency). Data are shown as mean ± s.e.m. (n = 5 mice per group, two-way ANOVA with Tukey’s multiple comparison test). (g) Contextual fear conditioning test schematic and timeline. (h) Quantification of freezing time during fear recall, and in a new context. Data are mean ± s.e.m. (n = 5 mice/group, one-way ANOVA with Tukey’s multiple comparison test). (i) Representative M2 immunofluorescence 1 week after treatment: NeuN (neurons), Iba1 (microglia), GFAP (astrocytes) and DAPI (nuclei). Images share one scale bar. (j) Quantification of marker-positive percentage (NeuN⁺/Iba1⁺/GFAP⁺ of DAPI⁺, left column) and cell densities (right column). Data are shown as mean ± s.e.m. (n = 9 mice per group, one-way ANOVA with Tukey’s multiple comparison test).

We therefore tried to test whether Dual-NIR Switch could lead to interference with normal behaviors (Figure 6b). General locomotor activity level and anxiety were assessed by the open filed test followed by the rotarod test to evaluate motor coordination^42^. No statistically significant difference of behavioral readouts was observed across all treatment groups (Figure 6c–f and Figure S19a). Cognitive function, assessed by contextual fear conditioning^43^, also remained intact (Figure 6g,h and Figure S19b,c). These results confirm that the Dual-NIR Switch does not impair normal motor or cognitive ability.

In contrast to the high invasiveness of conventional optogenetics, Dual-NIR Switch induces negligible tissue damage. Histological analysis 1 week post-treatment showed no significant loss of neurons, and glial activation was minimal: microglia exhibited resting morphology, and GFAP⁺ astrocytes were sparse near the injection site^44^ (Figure 6i,j). Tissue integrity was similarly preserved at 4 weeks (Figure S19d,e), indicating preserved tissue integrity and the absence of notable acute or chronic inflammation.

A key safety advantage of Dual-NIR Switch is its decoupling of neuronal activation from sustained NIR exposure. By limiting NIR delivery to brief initiating and terminating illuminations, it substantially reduces cumulative thermal load. Thermal imaging confirmed that the sequential NIR illumination at 980 nm followed by 808 nm significantly lower temperature rises compared to continuous-wave or pulsed illumination regimes used in conventional approaches (Figure S20). Under all behavioral stimulation patterns used in this study, including the highest intensity (50 mW/mm²), temperature increases remained within a safe, transient range (peak ≤ 40.1 °C; Figure S21). Thus, by decoupling NIR exposure from neuronal activation, Dual-NIR Switch substantially mitigates photothermal burden and associated tissue damage risk.

Taken together, Dual-NIR Switch exhibits an excellent safety profile, including negligible thermal effects, preserving normal behavior, and causing no detectable tissue damage or inflammatory response. All results support the potential of Dual-NIR Switch for long-term in vivo applications.

## Conclusions

We have developed Dual-NIR Switch, a wireless neuromodulation technology that provides temporally precise and duration-tunable control of neuronal activity in freely moving mice. Its core component, the engineered odUCNPs emits blue and green light under 980 nm and 808 nm excitation, respectively, perfectly matching the activation and inactivation spectra of the step-function opsin SOUL. The combination of odUCNPs with SOUL allows neuronal activation to be fully decoupled from sustained NIR delivery: a brief 980 nm illumination initiates excitation that persists without further energy input, and a subsequent 808 nm illumination rapidly terminates it. Leveraging the deep-tissue penetration of NIR light, Dual-NIR Switch supports transcranial modulation at depths of ∼6 mm, enabling truly remote, tether-free operation.

We successfully demonstrated that the technology could control behavior across a broad spectrum of durations, including short-duration locomotion, medium-duration feeding suppression, and long-duration reward processing. Moreover, all components exhibit excellent biocompatibility, with no detectable impairment of motor or cognitive function and no significant tissue damage or inflammatory response, even after long-term use.

Looking forward, Dual-NIR Switch can be further refined. The upconversion efficiency of odUCNPs could be enhanced by boosting excitation harvesting^45–49^, or by further suppressing surface quenching^50^. This improvement would enable Dual-NIR Switch to operate at lower NIR power levels, thereby increasing its sensitivity and achieving higher temporal resolution. Furthermore, by functionalizing the nanoparticle surface to cross the blood-brain barrier^51,52^, the entire Dual-NIR Switch system could be delivered via intravenous injection and operated noninvasively by transcranial NIR illumination.

In summary, Dual-NIR Switch is an encouraging step toward optical wireless neuromodulation, with the advantages of tunable duration control, minimal invasiveness, and high biocompatibility. Beyond its utility for neuroscience research in behaving animals, the ability to drive sustained yet reversible neuronal activation opens promising avenues for therapeutic applications in various neurological disorders.

## Author Contributions

Z.X. and L.P. contributed equally. Z.X., L.P., and J.L. conceived the project and designed the experiments. Z.X. conducted the synthesis and characterization of the materials. J.D., P.X. and Y.C. provided theoretical guidance and experimental design support for the patch-clamp experiments. Z.X., L.P., P.X., S.S. and X.Y. performed the cell culture and patch-clamp experiments. Z.X., L.P., B.H., and Y.H. carried out the in vivo electrophysiological recording in mice. Z.X. and Y.H. conducted behavioral experiments. Z.X., L.P., Y.H., B.H., and P.X. participated in data analysis. Z.X., Y.W., P.G. and X.Y. conducted the immunostaining and biocompatibility assessments. Z.X. completed the manuscript draft. J.L., L.P., J.D. and P.X. revised the manuscript. J.L. and L.P. supervised the entire project.

## Supporting Information

Supporting information includes materials & methods, figures, tables, video list and references.

## Competing interest notes

The authors declare no competing interest.

## Acknowledgements

This work was supported by STI2030-Major Projects (2021ZD0202200 and 2021ZD0202203), National Natural Science Foundation of China (32271144), the Chinese Academy of Sciences Project for Young Scientists in Basic Research (YSBR-139), and Lingang Laboratory Grant LG-QS-202202-03. The authors thank Dr. Danqian Liu at CEBSIT for her critical comments on this work. The authors would also acknowledge Mr. Lijun Pan of the Electron Microscopy Core Facility at CEBSIT for assistance with TEM and SEM sample preparation, Mr. Zhengyu Lu and Mrs. Qianru Ma of the Optical Imaging Core Facility at CEBSIT for support with confocal imaging, and the Mice Facility at CEBSIT for assistance with behavioral assays.

