## Supporting Information for "An orthogonal near-infrared optical switch for wireless neuromodulation in freely behaving mice"

### Materials and Methods

#### Materials

All chemicals used in this paper were used without further purification. Oleic acid (OA, 90%), 1-octadecene (ODE, 90%), ammonium fluoride ($NH_4F$, ≥98%), $YCl_3·6H_2O$ (99.99%), $YbCl_3·6H_2O$ (99.9%), $NdCl_3·6H_2O$ (99.9%), $ErCl_3·6H_2O$ (99.9%), $TmCl_3 · 6H_2O$ (99.99%), bovine serum albumin (BSA, ≥98%), L-glutathione reduced (GSH, ≥98.0%), sodium chloride (NaCl, ≥99%), potassium chloride (KCl, ≥99.0%), calcium chloride dihydrate ($CaCl_2·2H_2O$, ≥96.0%), HEPES (≥99%), D-(+)-Glucose (≥99.5%), potassium hydroxide (KOH, 90%), EGTA (≥97.0%), phosphocreatine di(tris) salt (≥97%), potassium gluconate (K-gluconate, 97.0-103.0%), GTP sodium salt hydrate (≥95%), ATP magnesium salt (≥95%), Triton X-100 were purchased from Sigma-Aldrich. Sodium trifluoroacetate (NaTFA, ≥98%) was purchased from TCI. Methanol (≥99.5%), sodium hydroxide (NaOH, ≥96.0%), cyclohexane (≥99.5%), anhydrous ethanol (≥99.7%) were purchased from Sinopharm Chemical Reagent Co., Ltd. DMEM, fetal bovine serum (FBS) and PBS were from Gibco (Invitrogen). ExFect transfection reagent was purchased from Vazyme. All antibodies were purchased from Abcam. Primary antibody includes Iba11 (ab178847), NeuN (ab208942), GFAP (ab68428) and c-Fos (ab208942). Secondary antibody includes donkey anti-rabbit IgG H&L (ab175470) and goat anti-mouse IgG H&L (ab150113).

#### Synthesis of odUCNPs

The multilayer core-shell UCNPs were synthesized using a thermal decomposition method. To achieve orthogonal emissions, a layer-by-layer epitaxial growth approach was utilized to incorporate distinct lanthanide dopants into specific shell layers at precisely controlled concentrations[1–3].

1. **Synthesis of core nanoparticles: $NaYF_4$: 0.5 mol% $Nd^{3+}$, 20 mol% $Yb^{3+}$, 2 mol% $Er^{3+}$**

Rare earth chlorides, including $YCl_3\cdot6H_2O$ (0.775 mmol), $NdCl_3\cdot6H_2O$ (0.005 mmol), $YbCl_3\cdot6H_2O$ (0.20 mmol), and $ErCl_3\cdot6H_2O$ (0.02 mmol) were weighed and introduced into a 50 mL three-neck flask. To the flask, 6 mL of OA and 15 mL of ODE were added. Next, under an argon atmosphere, the mixture was heated to 165 °C for 40 minutes to remove water. Then, the system was allowed to cool down to 50 °C. Subsequently, 10 mL of methanol containing NaOH (2.5 mmol) and $NH_4F$ (4 mmol) were carefully transferred to the flask and stirred at 40 °C for 1 hour. Methanol was then removed by heating the system to 90 °C under vacuum for complete removal. Finally, under argon flow, the temperature was raised to 290 °C and maintained for 1 hour and 30 minutes.

After cooling down to room temperature, 10 mL of anhydrous ethanol was added, and the mixture was centrifuged at 10,000 rpm for 5 minutes. Then, the supernatant was discarded, and the precipitate was dispersed in 10 mL of cyclohexane by ultrasound. Repeat the washing steps twice. Finally, the precipitate was dispersed in 10 mL of cyclohexane for further use.

#### 2. Synthesis of core-shell ($CS_1$) nanoparticles: core@β-$NaYF_4$: 20 mol% $Nd^{3+}$

NaTFA (1.25 mmol) was dissolved in oleic acid (2.5 mL) with heating under argon. After the mixture (NaTFA-OA) was cooled down to room temperature. Rare earth chlorides, including $YCl_3\cdot6H_2O$ (0.80 mmol) and $NdCl_3\cdot6H_2O$ (0.20 mmol) were weighed and introduced into a 50 mL three-neck flask. To the flask, 10 mL of OA and 20 mL of ODE were added. Next, under an argon atmosphere, the mixture was heated to 165 °C for 40 minutes to remove water. Then, the system was allowed to cool down to room temperature following the step that 2.5 mL of NaTFA-OA solution was added to the flask and stirred for 10 min. Afterward, the pre-prepared core-cyclohexane solution was added dropwise at 60 °C. Cyclohexane was then removed by heating the system to 120 °C under vacuum for complete removal. Finally, under argon flow, the temperature was slowly raised to 280 °C and maintained for 1 hour. Washing was performed following the same procedure. Finally, the precipitate was dispersed in 10 mL of cyclohexane for further use.

**3. Synthesis of core-shell-shell ($CS_2$) nanoparticles: $CS_1$@β-$NaYF_4$**

Rare earth chlorides, $YCl_3 \cdot 6H_2O$ (1.00 mmol) were weighed and introduced into a 50 mL three-neck flask. The other process for coating the second shell was same as the above-mentioned core-shell ($CS_1$) nanoparticles synthesis.

**4. Synthesis of core-shell-shell-shell ($CS_3$) nanoparticles: $CS_2$@β-$NaYF_4$: 20 mol% $Yb^{3+}$, 0.5 mol% $Tm^{3+}$**

Rare earth chlorides including $YCl_3 \cdot 6H_2O$ (0.795 mmol), $YbCl_3 \cdot 6H_2O$ (0.20 mmol), $TmCl_3 \cdot 6H_2O$ (0.005 mmol), were weighed and introduced into a 50 mL three-neck flask. The other process for coating the third shell was same as the above-mentioned core-shell ($CS_1$) nanoparticles synthesis.

**5. Synthesis of core-shell-shell-shell ($CS_4$) nanoparticles: $CS_3$@β-$NaYF_4$**

Rare earth chlorides, $YCl_3 \cdot 6H_2O$ (1.00 mmol) were weighed and introduced into a 50 mL three-neck flask. The other process for coating the fourth shell was same as the above-mentioned core-shell ($CS_1$) nanoparticles synthesis.

**Surface modification of odUCNPs with glutathione**

UCNPs surface was modified with GSH through ligand exchange[4]. GSH solution (0.125 mmol in 10 mL $H_2O$, pH adjusted to ~1 with HCl) was mixed with 2 mL UCNPs dispersed in cyclohexane. The biphasic mixture was stirred for more than 60 min. Then, the aqueous phase was collected, centrifuged (13,000 rpm, 6 min), and the pellet was washed with ethanol by repeated centrifugation. The final GSH-modified UCNPs were dispersed in 1 mL of ultrapure water for further biological use.

**Characterization of nanoparticles**

UCNPs were characterized by JEM-2100F (JEOL, Japan) at accelerating voltage of 200 kV for morphology, crystal structure (HRTEM), and elemental analysis (EDS/HAADF-STEM). The particle size distribution was determined by Zetasizer Advance (Malvern Panalytical, UK). Upconversion luminescence spectra were obtained on an FS5

Spectrofluorometer (Edinburgh Instruments, UK) coupled with 980 nm and 808 nm lasers (Changchun New Industries Optoelectronics Technology Co., Ltd., China) as the excitation sources.

**Cell culture and transfection**

HEK293T cell lines were purchased from the Cell Bank of Type Culture Collection of the Shanghai Institute of Cell Biology, Chinese Academy of Sciences. Cells were grown in monolayers in DMEM supplemented with 10% heat-inactivated FBS at 37 °C under a humidified 5% CO2 atmosphere. HEK293T cells in 24-well plates (0.3-1×10^5 cells/well) were transfected with the target plasmid using a commercial reagent. The transfection complex, prepared by mixing plasmid and reagent (6 μg plasmid pre-diluted in 300 μL opti-MEM) and incubating for 15 min, was added (50 μL/well). After 8 h, the medium was replaced with fresh DMEM, and cells were cultured for 24-72 h before use.

**Scanning electron microscopy of HEK293T cells**

SEM samples of HEK293T cells were prepared by sequential processing. Cells were rinsed with phosphate buffer (PB, 0.1 M, pH 7.4), then fixed with 2.5% glutaraldehyde/PB (1 h at RT), and next washed with cold PB three times at 4°C (10 minutes per wash). After post-fixation with 1% $OsO_4$/PB (1 h at 4°C, dark) and repeated washes with cold PB, dehydration was carried out in a graded ethanol series (30%, 50%, 70%, 80%, 90%, 100%; 10 min per step). Samples underwent 4-h critical-point drying by EM CPD 300 (Leica Microsystems, Germany), were mounted on conductive adhesive, sputter-coated with ~10 nm gold, and examined under a Gemini SEM 300 (ZIESS, Germany).

**Whole-cell patch-clamp recording of HEK293T cells**

Whole-cell patch-clamp recordings were obtained from HEK293T cells using borosilicate glass pipettes (BF100-58-10; Sutter Instrument, USA) pulled on a P-97 micropipette puller (Sutter Instrument, USA). Pipette resistances were 15–20 MΩ. The extracellular solution contained (in mM): 134 NaCl, 2.9 KCl, 4 $CaCl_2 \cdot 2H_2O$, 10 HEPES, and 10 D-glucose (pH

7.81, 292 mOsmol/L). The internal (pipette) solution contained (in mM): 100 K-gluconate, 10 HEPES, 10 EGTA, 2 $CaCl_2 \cdot 2H_2O$, 10 KCl, 2 Mg-ATP, 0.3 Na-GTP, and 2 phosphocreatine di(tris) salt (pH 7.41, 271 mOsmol/L). NIR lasers (980 & 808 nm; Changchun New Industries Optoelectronics Technology Co., Ltd., China) were coupled via mirrors into an BX51W microscope (Olympus, Japan) for delivery of irradiation through the objective lens to the field of view below.

HEK293T cells were incubated with 400 μL of GSH-modified UCNPs (in PBS) for 15 minutes. After removal of the medium and a gentle rinse with extracellular solution (ES), the dish was filled with fresh ES for recording. Under IR-1000E camera guidance (Dage-MTI, USA), a patch pipette filled with internal solution (IS) was positioned onto a cell using an MPC-200 micromanipulator (Sutter Instrument, USA) to form a giga-ohm seal. Gentle negative pressure was applied to establish the whole-cell configuration. Signals were amplified by a MultiClamp 700B amplifier, digitized with an Axon Digidata 1550A (Molecular Devices, USA), and sampled at 10 kHz.

**Virus and odUCNPs injections**

AAV9-hSyn-SOUL-P2A-tdTomato (titer: 5.5E+12 GC/mL): viral vectors were purchased from Addgene. Viral packaging was performed by Gene Editing Core Facility of CEBSIT, Chinese Academy of Sciences. AAV9 -CaMKIIa-SOUL-mCherry (titer: 1E+13 GC/mL) were purchased from PackGene Biotech (Guangzhou, China). Surface modified odUCNPs were centrifuged and redispersed in aCSF. Both the viral solution and the nanomaterials were loaded using glass pipette with micro-syringes.

Mice anesthetized with isoflurane were fixed in a stereotaxic frame (RWD Life Science, China) with body temperature maintained by a heating pad. After exposing the skull and identifying Bregma/Lambda, pipettes were lowered to the following coordinates: M2 (+1.00 mm AP, ±0.50 mm ML, -0.50 mm DV), LHA (-1.41 mm AP, ±1.15 mm ML, -0.55 mm DV), VTA (-3.08 mm AP, +0.40 mm ML, -4.55 mm DV) through a small craniotomy. Virus (300 nL/site) and odUCNPs (1000 nL/site) were injected at 20 nL/min and 15 nL/min, respectively. The pipette was kept in place for 15 min post-injection before withdrawal. At last, wound closure was conducted using surgical suture or 3M Vetbond Tissue Adhesive.

**Bio-transmission electron microscope of brain tissue**

Mice received a stereotaxic injection of odUCNPs-aCSF (1000 nL) or aCSF (1000 nL) into the M2 region and were perfused at specified times thereafter. Tissue samples were primarily fixed with 2.5% glutaraldehyde in PB (0.1 M, pH 7.4) at 4°C for 24 h, followed by post-fixation with 1% $OsO_4$/PB (2 h at RT). After dehydration in a graded ethanol series (30%, 50%, 70%, 90%, 100%; 15 min per step), samples were infiltrated with Epon 812 resin using propylene oxide as a transition solvent, and polymerized at 60°C for 48 h. Ultrathin sections (70 nm) were cut with a Leica Ultracut microtome, placed on 50-mesh copper grids, double-stained with 2% uranyl acetate and lead citrate. Finally, the grids were imaged using a JEM-1230 (JEOL, Japan) TEM at accelerating voltage of 80 kV.

**In vivo multichannel electrophysiology recording of M2 in mice**

*In vivo* electrophysiological recordings were conducted 1-3 days after odUCNPs injection. Mice anesthetized with isoflurane were fixed in a stereotaxic frame (RWD Life Science, China) with body temperature maintained by a heating pad. A craniotomy was performed over the target brain region followed by removal of the skull and dura. A 16-channel tetrode (Kedou Brain-Computer Technology Co., Ltd., China) was lowered 0.4–0.6 mm below the cortical surface. Neuronal signals were acquired, amplified, and digitized using a Cerebus recording system with a CerePlex μ headstage (Blackrock Microsystems, USA), sampled at 1 kS/s. Spikes were bandpass-filtered between 250 Hz and 750 Hz. The acquired data was spike-sorted offline using Blackrock Offline Spike Sorter. Sorted units were subsequently analyzed in MATLAB (Mathworks, USA) to extract and plot waveform shapes, generate raster plots, and compute firing rates. Onset and offset estimation of in vivo electrophysiology recordings: baseline activity was defined as the interval from 0.5 to

29.5 s, and the baseline mean firing rate ($\mu_{base}$) was computed from this period. The peak firing rate ($FR_{peak}$) was identified from the entire time series. A peak-relative threshold was then defined as $\mu_{base} + 0.46 * (FR_{peak} - \mu_{base})$. Onset was defined as the first time bin preceding the peak in which the firing rate exceeded this threshold for at least three consecutive bins. Offset was defined as the first-time bin following the peak at which the firing rate fell below this threshold for at least three consecutive bins. The decay threshold (0.46 of the peak response relative to baseline) was chosen as an approximation to the half-maximum level that ensured a stable identification of the main response epoch in the presence of minor post-peak fluctuations.

**Immunostaining**

Following experimental treatments, mice were transcardially perfused with PBS and 4% PFA. The harvested brains were post-fixed in 4% PFA at 4°C overnight, then cryoprotected in sequential 15% and 30% sucrose (in PBS) until equilibrated. Tissues were embedded in OCT, frozen, and coronally sectioned at 30 μm thickness using a CryoStar NX50 cryostat (Thermo Scientific, USA). Sections were blocked for 1 h at room temperature with 10% BSA and 0.3% Triton X-100 in PBS, then incubated with primary antibody (1:200 dilution in 3% BSA, 0.3% Triton X-100/PBS) overnight at 4°C. After five 5-min PBS washes, sections were incubated with fluorophore-conjugated secondary antibody (1:500 in 0.3% Triton X-100/PBS) for 2 h at room temperature, washed three times with PBS, counterstained with DAPI (Solarbio, China) for 10 min, washed again five times, and finally mounted with Fluoromount-G (Southern Biotech, USA) and stored at 4 °C in the dark until further imaging.

**Fluorescence imaging of cells and brain slices**

Cellular imaging was conducted using a FV3000 confocal microscope (Olympus, Japan). For brain slices, high-magnification images were acquired with a C2 confocal microscope (Nikon, Japan), while overview, low-magnification images were captured using a VS120 slide scanner (Olympus, Japan). Imaging parameters were kept consistent across all samples within the same experiment to ensure comparability.

**Thermal effect evaluation**

After cranial hair removal and anesthesia, mice were head-fixed in a stereotaxic frame (RWD Life Science, China). A FOTRIC 692 thermal imaging camera (FOTRIC, China) and the NIR light source were positioned above the head, with the irradiation area adjusted via real-time thermal feedback. Each mouse received eight sequential NIR stimulation protocols (all pulses at 5 Hz, 0.1 s width unless stated): 1) 980 nm, 80 mW/mm$^2$, CW, 30 s; 2) 980 nm, 80 mW/mm$^2$, pulsed, 30 s; 3) 808 nm, 80 mW/mm$^2$, CW, 30 s; 4) 808 nm, 80 mW/mm$^2$, pulsed, 30 s; 5) 980 nm → 808 nm (80 mW/mm$^2$ each, CW): 5 s → 20 s interval → 5 s; 6) 980 nm → 808 nm (40 mW/mm$^2$ each, CW): 5 s → 60 s interval → 5 s; 7) 980 nm → 808 nm (50 mW/mm$^2$ each, CW): 10 s → 60 s interval → 30 s; 8) 980 nm → 808 nm (60 mW/mm$^2$ each, pulsed): 30 s → 60 s interval → 30 s.

**Locomotion enhancement test related to M2**

Four weeks after unilateral AAV9-CaMKIIa-SOUL-mCherry injection into the M2 cortex (+1.00 mm AP, +0.50 mm ML, -0.50 mm DV), odUCNPs were injected one day before behavioral testing at the same site. Mice were handled for 3 consecutive days and acclimatized to the testing room (100 lux, 23 °C, 50% humidity) for 1 h prior to the experiment. The test was conducted in a square open-field arena (16×16×20 cm). After a 3-min habituation, mouse behavior was recorded for 60 s. During recording, two 5-s NIR illuminations (40 mW/mm$^2$, CW) were delivered: 980 nm at 15 s and 808 nm at 40 s. For proof-of-principle validation, NIR was delivered via an optical fiber coupled with a collimator and controlled manually, a procedure consistent across all behavioral experiments. Behavioral data (velocity, distance moved, heat maps, angular velocity, rotations, mobility) were analyzed using EthoVision XT 10 (Noldus). Locomotor trajectories for Pre-, During-, and Post-period were plotted in MATLAB using exported positional data. For circling analysis, a full circle as a 180° counterclockwise rotation (around the body or arena center) and assigned positive values to counterclockwise angular velocity.

**Free-access feeding test related to LHA**

Four weeks after bilateral AAV9-CaMKIIa-SOUL-mCherry injection into the LHA (-1.41 mm AP, ±1.15 mm ML, -5.35 mm DV), odUCNPs were bilaterally injected prior to behavioral testing at the same sites. Mice were handled for 3 consecutive days, food-deprived 24 h before the test. Before starting the experiments, mice were acclimatized to the testing room (50 lux, 23 °C, 50% humidity) for 1 h. Weight of mouse and a food-fille food bunk were measured first. After a 3-min free exploration in a square arena (40 × 40 × 40 cm), the food bunk was placed against the center of one wall[5]. Recording commenced when the mouse first entered a pre-defined "food zone" (ROI). During the 9-min 20-s trial, two 10-s NIR illuminations (50 mW/mm², CW) were delivered: 980 nm at 3 min and 808 nm at 6 min 10 s. After the test, the mouse was returned to its home cage with food restored, and consumption of food was quantified by re-weighing the bunk. Velocity, moved distance, and heat maps (Pre-/During-/Post-period) were analyzed using EthoVision XT 10 (Noldus).

**Conditioned place preference test related to VTA**

Four weeks after unilateral AAV9-hSyn-SOUL-tdTomato injection into the VTA (-3.08 mm AP, +0.40 mm ML, -4.55 mm DV), odUCNPs were unilaterally injected prior to testing into the same site. Mice were handled for 3 consecutive days. And before each session, mice were acclimatized to the testing room (30 lux, 23 °C, 50% humidity) for 1 h. The CPP test was performed in a three-compartment apparatus consisting of two large side compartments (20 × 20 × 20 cm each) connected by a narrower mid-compartment (20 × 7 × 20 cm). The compartments differed in visual and tactile cues. Guillotine doors between compartments could be opened or closed as needed[6].

Day 1 (Pre-test): After 5 min habituation in the closed central compartment, the mouse freely explored all compartments for 15 min while being recorded. Day 2 (Conditioning): Confined to one compartment (B), the mouse received three 30-min conditioning cycles, totaling 90 min. Each cycle consisted of 2 min habituation, a 30 s NIR illumination (980 nm, 60 mW/mm$^2$, 0.1 s-pulse, 5 Hz) at the start, and another 30 s illumination (808 nm, same parameters) at the end. Day 3 (Post-test): Identical to Day 1. Velocity, total distance moved, heat maps, and compartment occupancy were analyzed using EthoVision XT 10 (Noldus). Locomotor trajectories were plotted in MATLAB from exported positional data.

**Behavioral tests related to biocompatibility assessment**

Mice received different treatments (injections and/or NIR illumination) according to their experimental groups for three days prior to behavioral testing. During this period, mice were handled daily. One hour before each behavioral session mice were acclimatized to the testing room (30 lux, 23 °C, 50% humidity) for 1 h.

**Open Field Test:** Mice freely explored a square arena (40 × 40 × 40 cm) for 5 min (habituation) followed by a 15-min recorded test session. Velocity, moved distance, and center zone entries/duration were analyzed using EthoVision XT 10 (Noldus). Locomotor trajectories were plotted in MATLAB from exported positional data.

**Rotarod Test:** Training (one day prior) consisted of 5 trials: mice were placed on the rod accelerating from 10 to 80 rpm; trials ended upon falling or reaching max speed (30-min inter-trial intervals). The formal test included: 1) *Speed Test*: linear acceleration from 5 to 80 rpm over 60 s (3 trials; falling speed recorded); 2) *Duration Test*: constant 40 rpm for up to 200 s (3 trials; latency to fall recorded). All trials were separated by ≥30-min rest intervals.

**Contextual Fear Conditioning:** 1) *Fear Recall*: After 3-min habituation in the conditioning chamber (Med Associates Inc., USA), mice received five foot shocks (0.5 mA, 3 s, 60-s intervals) over 300 s, and remained in the chamber for 120 s. 24 h later, they were re-exposed to the same chamber for 300 s. 2) *New Context*: 24 h after fear recall, mice were placed in a modified chamber (smooth floor) for 3-min habituation and 300-s recording.

Freezing behavior was analyzed using VideoFreeze software (Med Associates; motion threshold: 18 pixels/frame; min freeze: 30 frames). Chambers were cleaned with odor-eliminator and 70% ethanol between trials.

**Animals**

All experimental procedures involving animals were approved by the CEBSIT, Chinese Academy of Sciences (NA-063-2024) and conducted in accordance with the NIH Guide for the Care and Use of Laboratory Animals. Male C57BL/6 mice (6-12 weeks old) were obtained from Shanghai Slac Laboratory Animal Co., Ltd. Before experiments, all mice were naïve and had not undergone any pharmacological treatment or surgical procedures. The animals were group-housed under standard specific pathogen-free (SPF) conditions with a 12/12-hour light/dark cycle, ambient temperature maintained at 23 ± 1°C, and relative humidity of approximately 35%. Food and water were provided *ad libitum*.

**Statistical analysis**

All data are presented as mean ± s.e.m. Statistical analyses were performed in GraphPad Prism 9.5. The specific statistical methods used for significance comparisons, including the tests for multiple comparisons, are provided in the figure legends for each panel. A standardized protocol was followed for the analysis of all behavioral data using EthoVision XT 10 (Noldus). Key parameters, including arena delineation, the animal detection threshold, and the criteria for initiating and ending analysis, were kept identical for all subjects within an experiment to eliminate inter-trial variability and human error. The data presented are the raw, unaltered outputs generated by the software.

### Supporting Figures

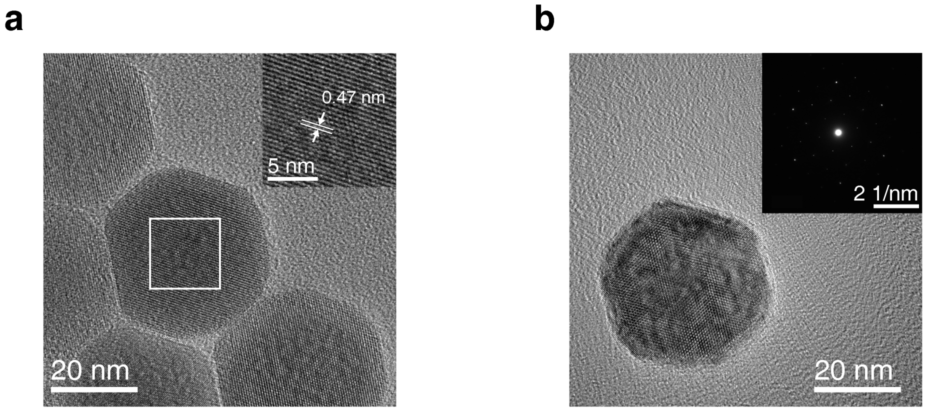

**Figure S1. High-resolution TEM images of odUCNPs.** (**a**) FFT patterns and (**b**) electron diffraction image.

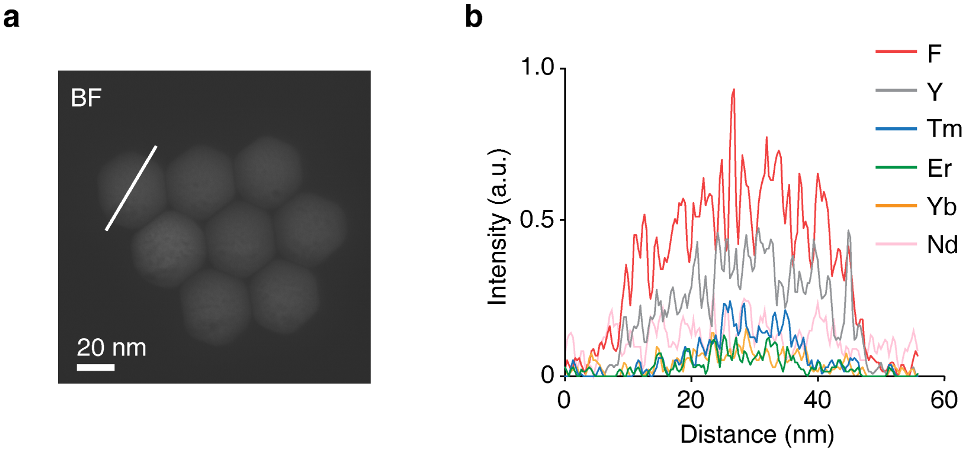

**Figure S2. Elemental characterization of odUCNPs.** (**a**) TEM image of odUCNPs. (**b**) Line-scanning of a single particle of odUCNPs. Data correspond to the white solid line in (a).

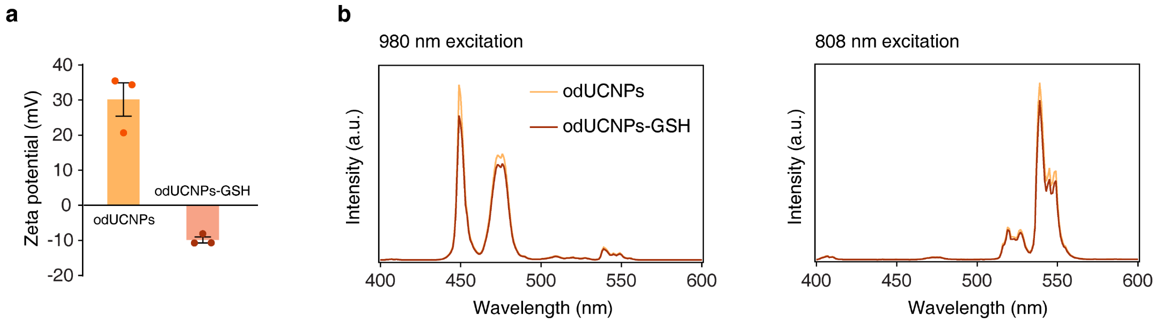

**Figure S3. Characterization of odUCNPs-GSH.** (**a**) Zeta potential of odUCNPs and GSH-modified odUCNPs. Data are shown as mean ± s.e.m. (n = 3 independent synthetic batches). (**b**) Emission spectra under 980 nm (left) and 808 nm (right) of odUCNPs and GSH-modified odUCNPs. GSH modification does not alter the emission peak wavelength and orthogonal emission.

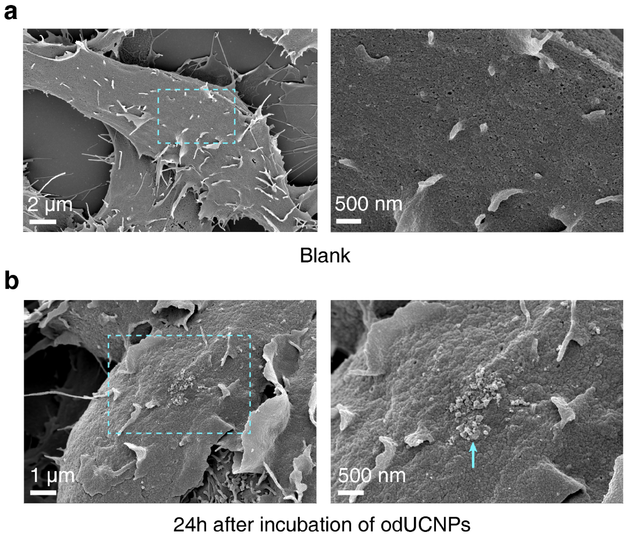

**Figure S4. Distributions of odUCNPs on 293T cells.** SEM images of 293T cells incubated without (**a**) or with odUCNPs (**b**) *in vitro*. Blue arrows indicate the interface of odUCNPs and 293T cells.

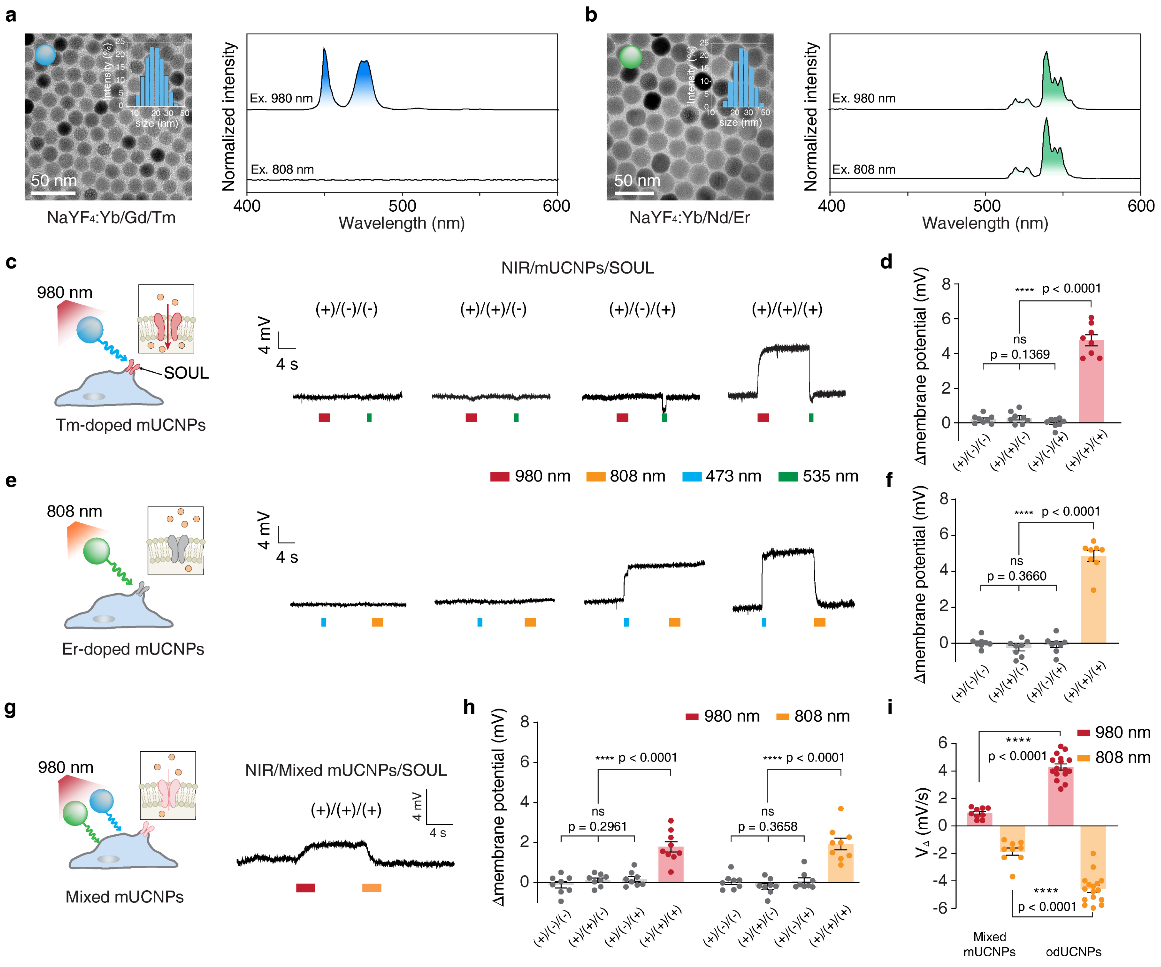

**Figure S5. mUCNPs modulate the membrane potential of SOUL-expressing 293T cells with low amplitude and slow kinetics.** (**a**) and (**b**) Structural and optical characterization of Tm- and Er-doped mUCNPs, showing TEM images with size distributions and normalized emission spectra under 980-nm and 808-nm excitation. (**c**) and (**e**) Whole-cell patch-clamp recordings in SOUL-expressing 293T cells incubated with Tm-doped (c) and Er-doped (e), with schematics and representative voltage traces under indicated conditions. (**d**) and (**f**) Quantification of NIR-illumination-induced membrane potential changes corresponding to (c) and (e), respectively. (**g)** Whole-cell patch-clamp recordings in SOUL-expressing 293T cells incubated with mUCNPs, with schematics and a representative voltage trace under (+)/(+)/(+) condition. Data are mean ± s.e.m. (n = 8 cells per group; one-way ANOVA with Tukey's multiple-comparison test). (**h)** Quantification of NIR-illumination-induced membrane potential changes corresponding to (g). Data are mean ± s.e.m. (n = 9 cells for experimental conditions and n = 8 cells for others; one-way ANOVA with Tukey's multiple-comparison test). (**i**) Quantification of velocity of membrane potential change for mUCNPs and odUCNPs. Data are shown as mean ± s.e.m. (n = 9 cells for mixed mUCNPs and n = 16 cells for odUCNPs, t-test).

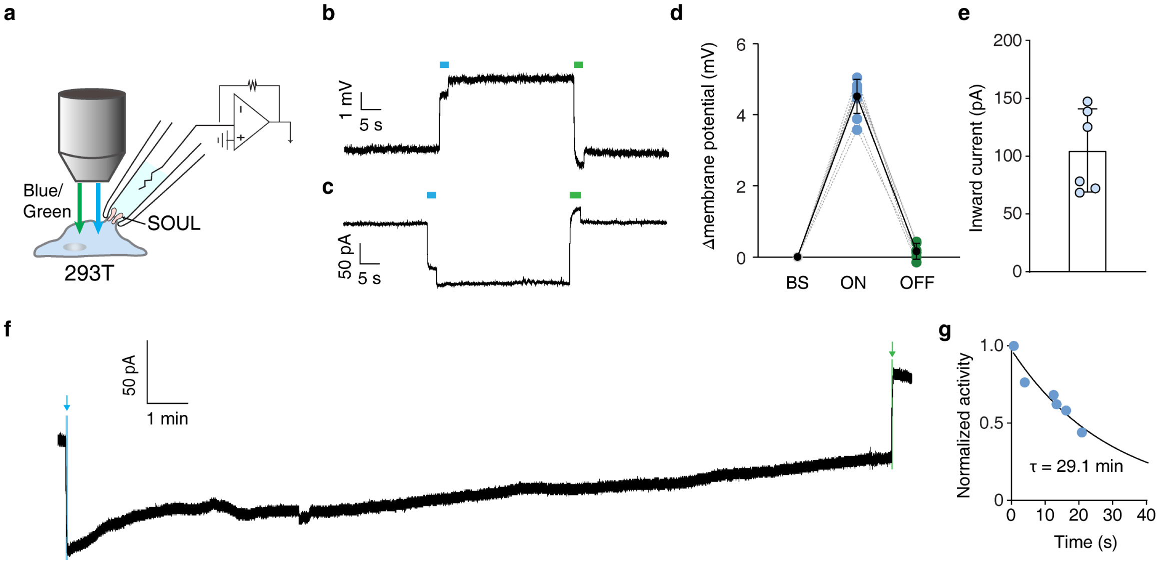

**Figure S6. Electrophysiological properties of SOUL-expressing 293T cells.** (**a**) Schematic of the patch-clamp setup to study electrophysiological properties of SOUL-expressing 293T cells upon sequential blue and green light illumination. (**b**) and (**c**) Representative voltage and current trace over time for a SOUL-expressing 293T cell *in vitro* upon blue-light activation (blue bar) and green light deactivation (green bar). (**d**) Membrane potential changes of individual 293T cell (dashed lines) and mean across cells (solid line, ± s.e.m.) during "ON" (35–45 s, after blue light illumination) and "OFF" (55–65 s, after green light illumination) compared to "BS" (5–15 s, before blue light illumination), (n = 6 cells). (**e**) Histograms of inward photocurrents of individual 293T cell. Data are shown as mean ± s.e.m. (n = 6 cells). (**f**) Representative long-duration activation current trace for a SOUL-expressing 293T cell. Blue and green shadows and arrows indicate the blue and green light illuminations. (**g**) Current-normalized activity over time with one phase exponential decay fit (solid line, time constant *tau* = 29.1 min).

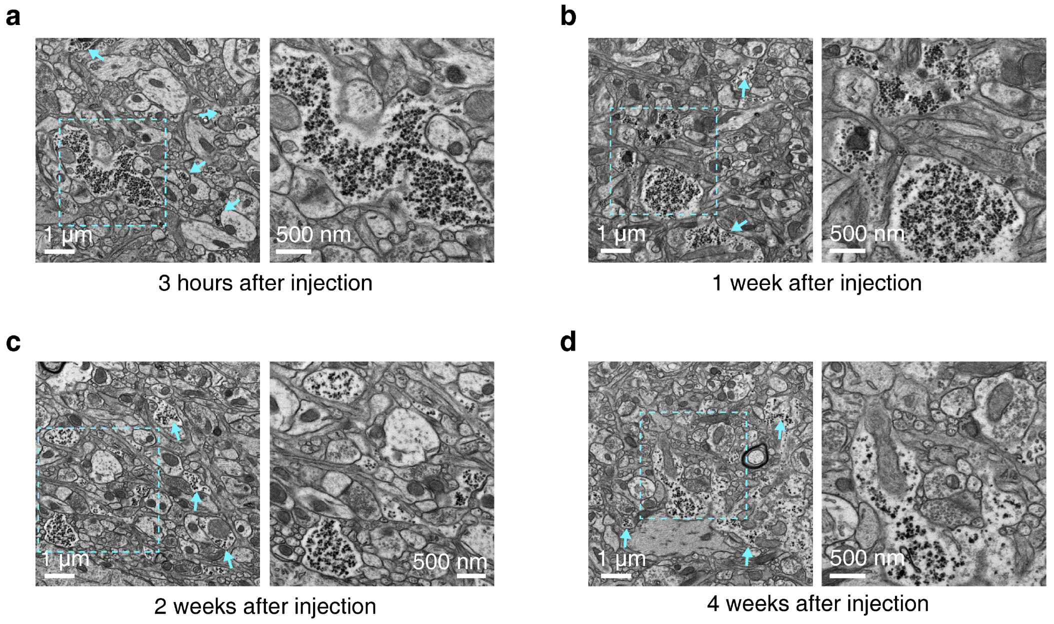

**Figure S7. Bio-TEM images of distributions of odUCNPs in M2 region over time. (a)–(d)** Blue arrows indicate the accumulation of odUCNPs. The Blue dashed outlines indicate the region of interest (ROI) shown in the zoom-in panel on the right.

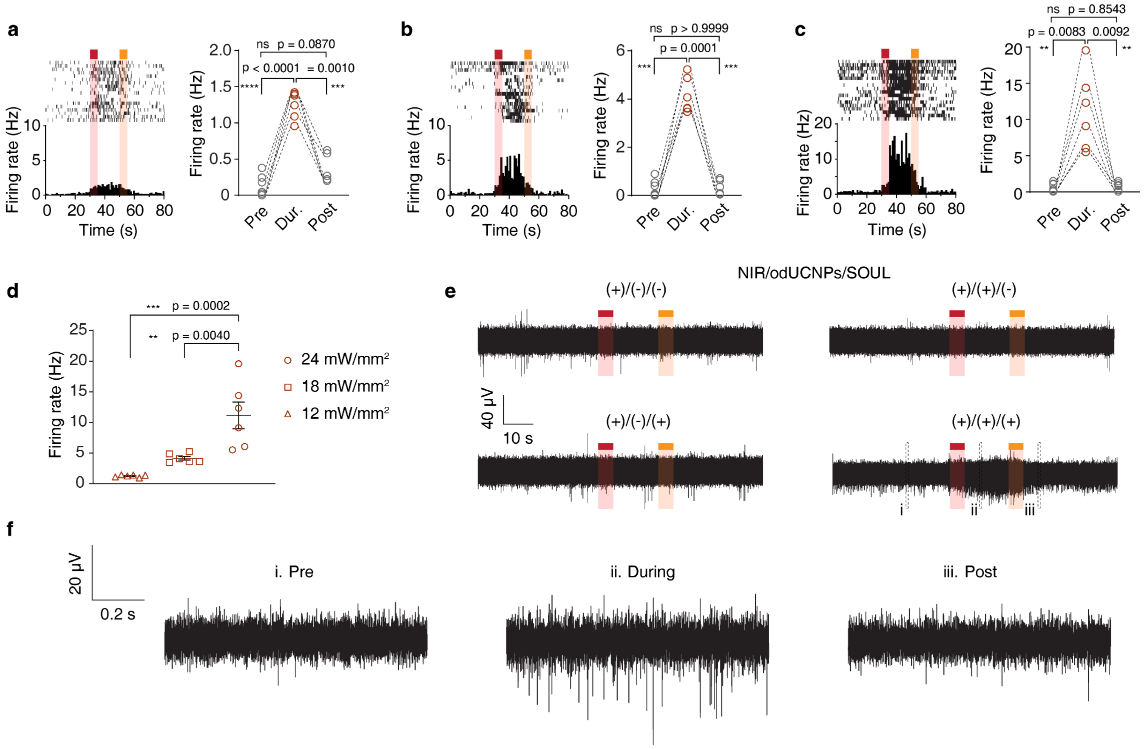

**Figure S8. Dual-NIR Switch induced firing properties of neuronal populations at**

**different NIR power density.** (**a**)–(**c**) Correspond to low (a), medium (b), and high power densities (c), respectively. Left, raster plots (top) and corresponding mean firing rate over time (bottom) under indicated conditions upon sequential NIR illumination at 980 nm (red) followed by 808 nm (orange). Right, quantification of the firing rate before 980 nm illumination (Pre, 0–15 s), after 808 nm illumination (Post, 65–80 s) and in between (During, 35–50 s). One-way ANOVA with Tukey's multiple comparison test. (**d**) Quantification of the firing rate in "During" period within different power densities. In (a)–(d), each point/point-set indicates a unit from one mouse averaged over 3 trials, n = 6 mice for each group, one-way ANOVA with Tukey's multiple comparison test. (**e**) Representative in vivo multichannel electrophysiological traces for the different conditions upon sequential NIR illumination at 980 nm (red) followed by 808 nm (orange). Traces from different time windows (dashed boxes) in the (+)/(+)/(+) group are magnified in (**f**). Both 980 nm and 808 nm were at a power density of 24 mW/mm², CW.

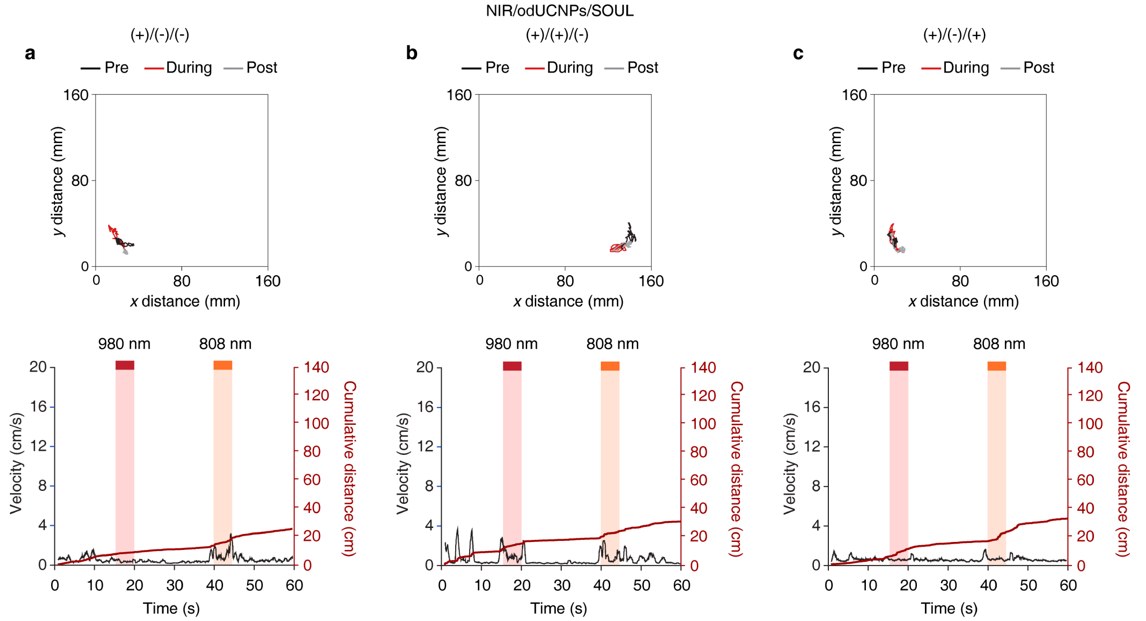

**Figure S9. Trajectories and dynamic of parameters of locomotion enhancement test related to M2 brain region in control group.** (**a**)–(**c**) Top panel, representative trajectory of the mouse before 980 nm illumination (Pre), in between (During) and after 808 nm illumination (Post). Bottom, kinetics of instant velocity (black) and cumulative distance moved (red) from a representative trail.

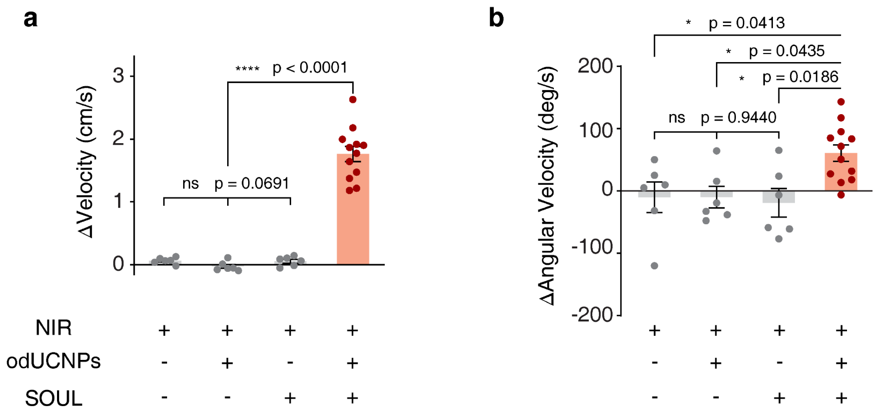

**Figure S10. Statistical analysis of locomotor parameters in the locomotion enhancement test related to M2 brain region for four groups.** Changes of velocity (**a**) and angular velocity (**b**) between "During" period and baseline (average of "Pre" and "Post"). Data are shown as mean ± s.e.m. (n = 12 mice for (+)/(+)/(+) and n = 6 mice for the other groups, each point is from 3 trails, one-way ANOVA with Tukey's multiple comparison test).

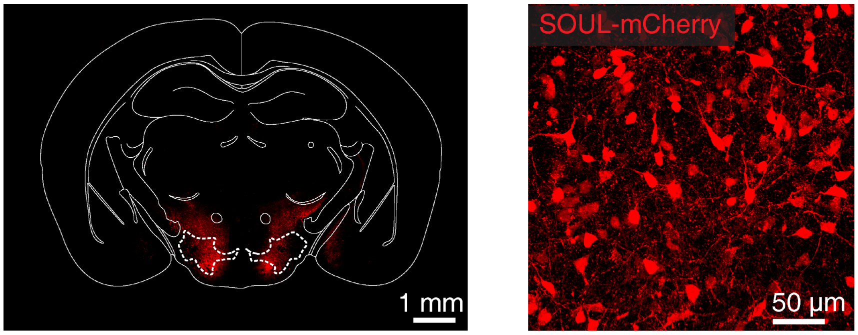

**Figure S11.** SOUL expression in the bilateral LHA. Left, Overview of SOUL expression. White dashed outlines indicate the LHA region. Right, Confocal image of the LHA.

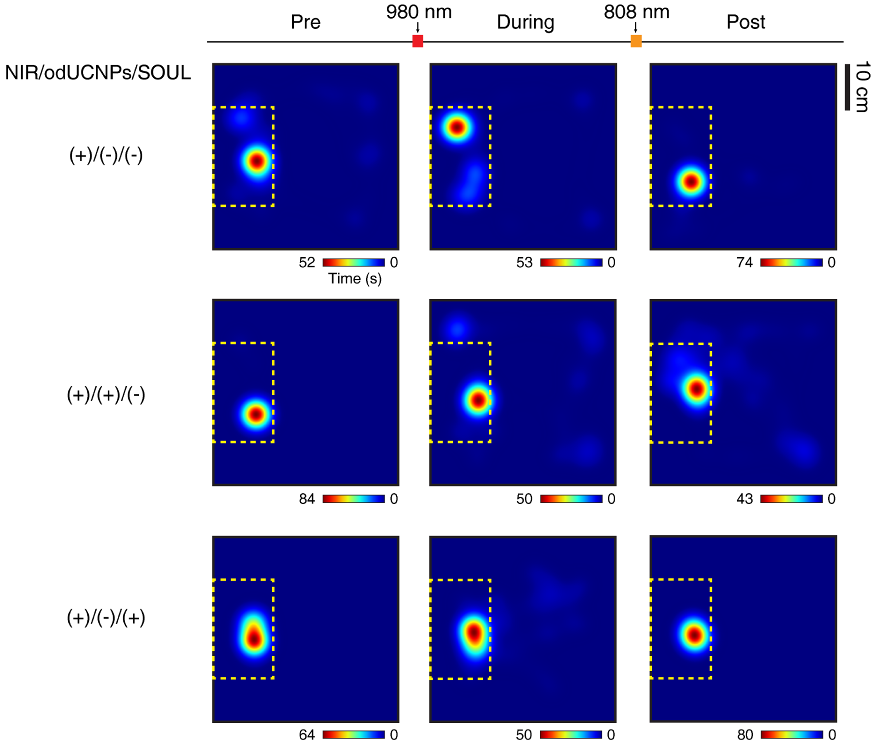

**Figure S12. Heatmaps of free-access feeding test related to LHA brain region in control group.** Representative heatmaps of “Pre”, “During” and “Post” period of each group. Scale bar is shown at the bottom right of each heatmap.

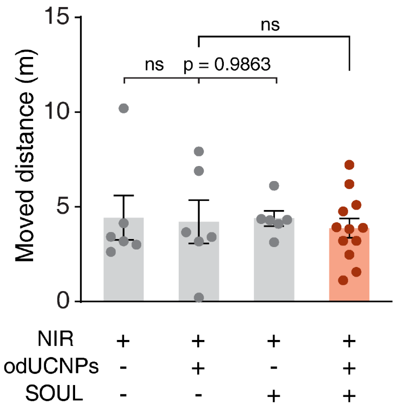

**Figure S13. Statistical analysis of locomotor parameters in the free-access feeding test related to LHA brain region for four groups.** Moved distance during the entire test for each group. Data are shown as mean ± s.e.m. (n = 12 mice for (+)/(+)/(+) and n = 6 mice for the other groups, one-way ANOVA with Tukey’s multiple comparison test, the full set p-values are provided in Supplementary Table 1).

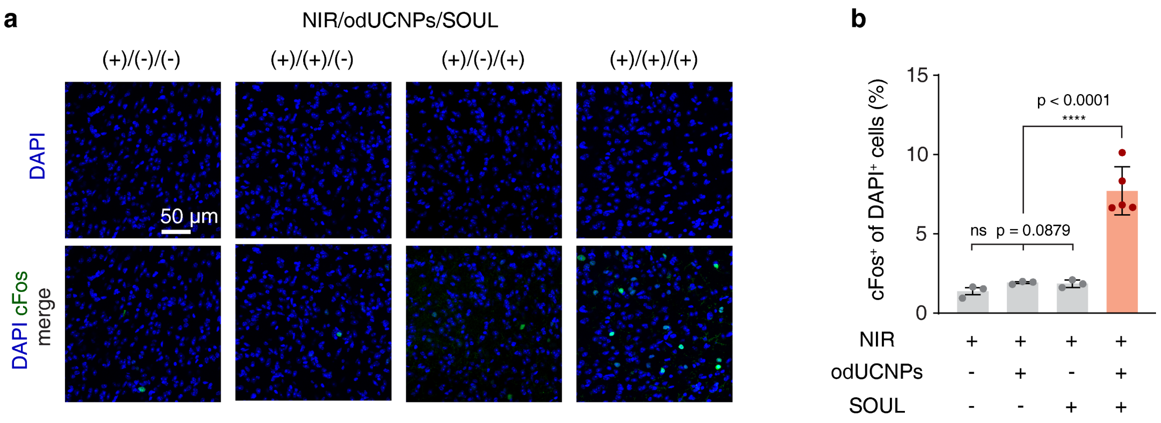

**Figure S14. Immunostaining of LHA region.** (**a**) Confocal images of c-Fos and DAPI in the LHA region under different experimental conditions. All images share one scale bar. (**b**) Statistical analysis of percentage of cFos+ cells within the DAPI+ cell population. Data are shown as mean ± s.e.m. (n = 5 mice for (+)/(+)/(+) and n = 3 mice for the other groups. Each point represents the average of fluorescence measurements from 3 brain slices. One-way ANOVA with Tukey's multiple comparison test).

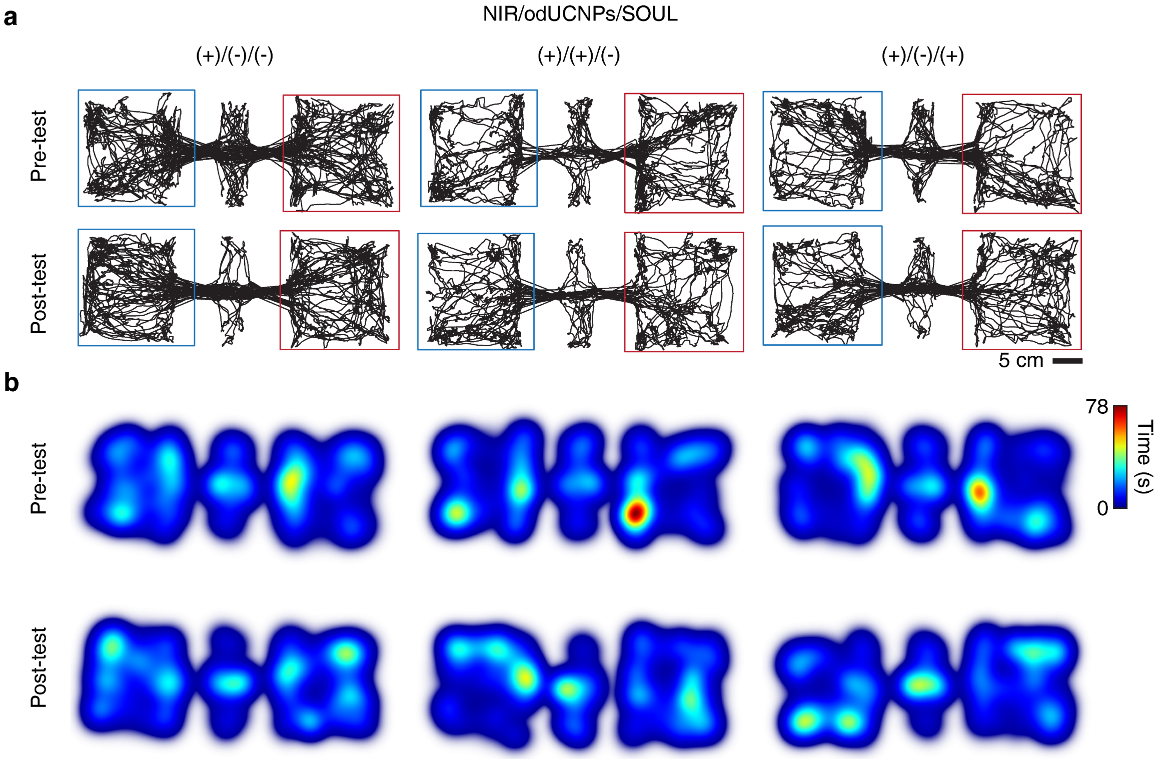

**Figure S15. Trajectories and heatmaps of CPP test related to VTA brain region for all**

**groups.** Representative trajectories (**a**) and corresponding heatmaps (**b**) in Pre- and Post-test for control groups. All heatmaps share one same scale bar.

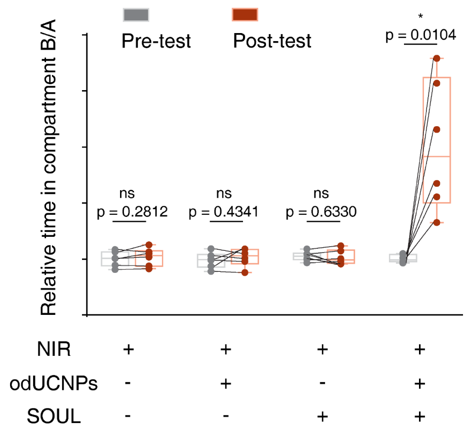

**Figure S16.** Statistical analysis of preference parameters in CPP test related to VTA brain region for four groups. The ratio of the time spent in compartment B to that in A. Data are shown as box plots indicating the min, max, and interquartile range, (n = 6 mice in each group, paired t-test).

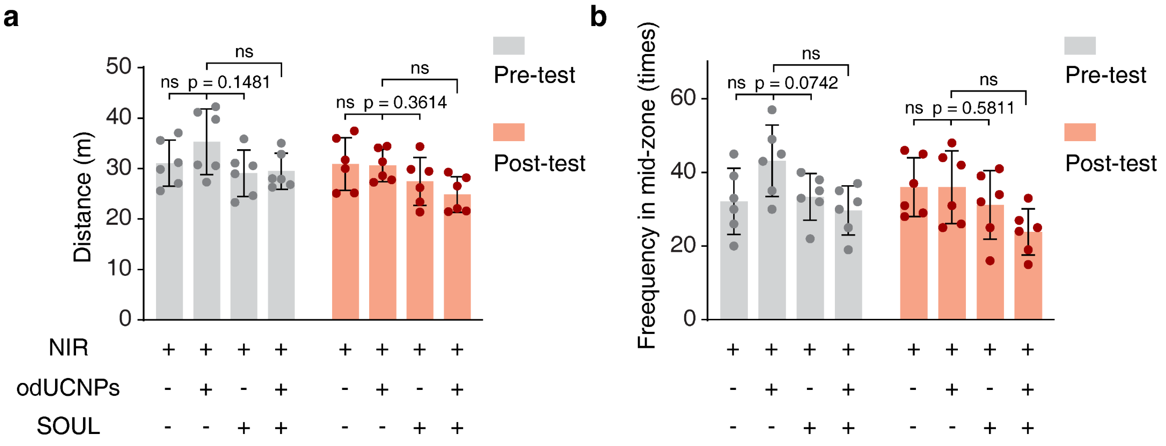

**Figure S17.** Statistical analysis of locomotor parameters in CPP test related to VTA brain region for four groups. Moved distance (**a**) and frequency in mid-zone (**b**) in Pre-test (grey points) and Post-test (red points). Data are shown as mean ± s.e.m. (n = 6 mice for each group, one-way ANOVA with Tukey's multiple comparison test, the full set p-values are provided in Supplementary Table 1).

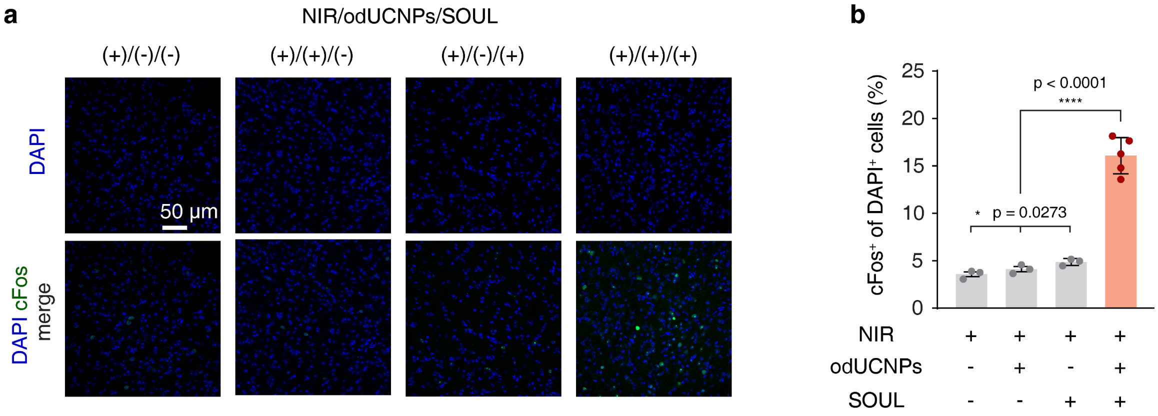

**Figure S18. Immunostaining of VTA region.** (**a**) Confocal images of c-Fos and DAPI in the VTA region under different experimental conditions. All images share one scale bar. (**b**) Statistical analysis of percentage of cFos+ cells within the DAPI+ cell population. Data are shown as mean ± s.e.m. (n = 5 mice for (+)/(+)/(+) and n = 3 mice for the other groups. Each point represents the average of fluorescence measurements from 3 brain slices. One-way ANOVA with Tukey's multiple comparison test).

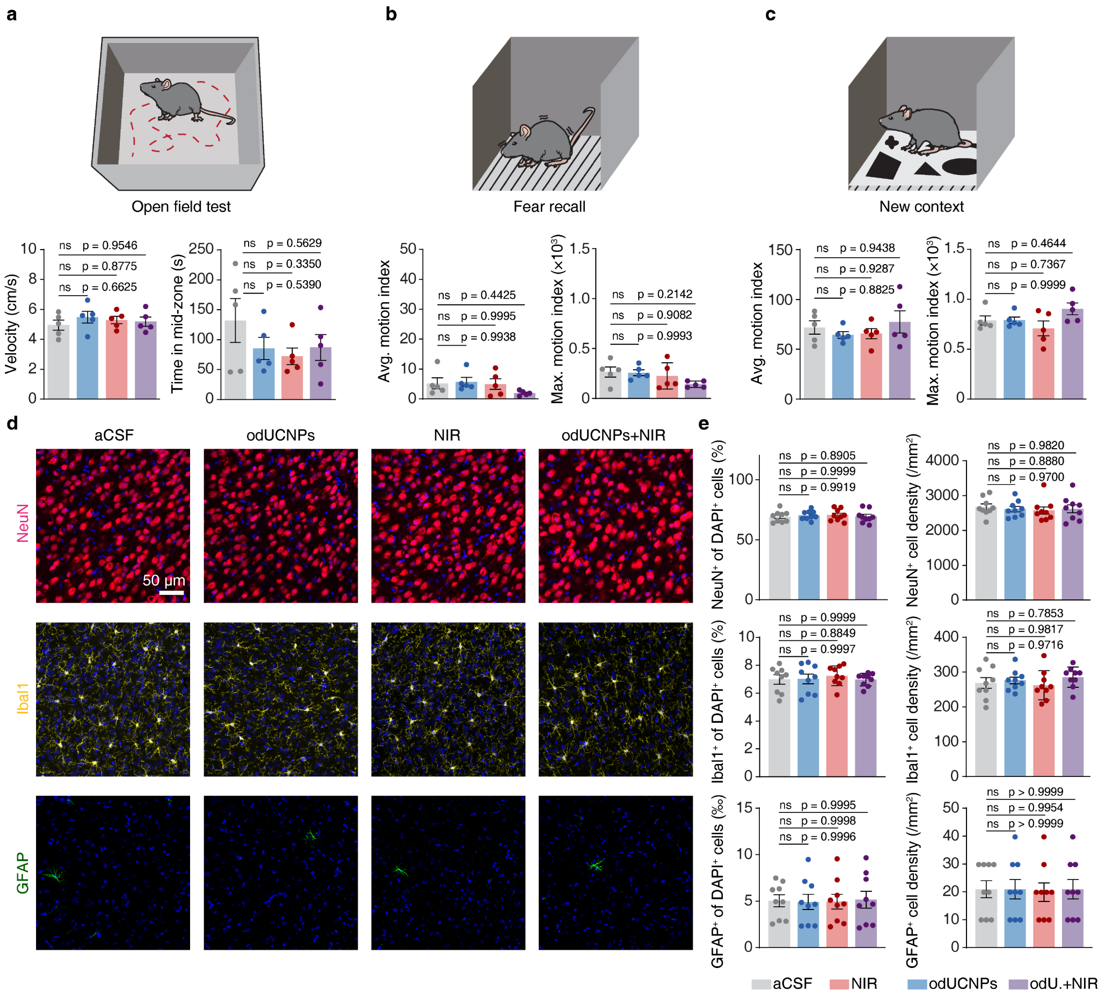

**Figure S19. Locomotor parameters and immunohistology in biocompatibility assessments of Dual-NIR Switch.** (**a**) Quantification of velocity and time in mid-zone in open field test. (**b**) Quantification of average motion index and maximum motion index in fear recall period of contextual fear conditioning test. (**c**) Quantification of average motion index and maximum motion index in new context period of contextual fear conditioning test. In (a) –(c), all data are shown as mean ± s.e.m. (n = 5 mice in each group, one-way ANOVA with Tukey's multiple comparison test). (**d**) Representative M2 immunofluorescence 4 weeks after treatment: NeuN (neurons), Iba1 (microglia), GFAP (astrocytes) and DAPI (nuclei). Images share one scale bar. (**e**) Quantification of marker-positive percentage (NeuN⁺/Iba1⁺/GFAP⁺ of DAPI⁺, left column) and cell densities (right column). Data are shown as mean ± s.e.m. (n = 9 mice per group, one-way ANOVA with Tukey's multiple comparison test).

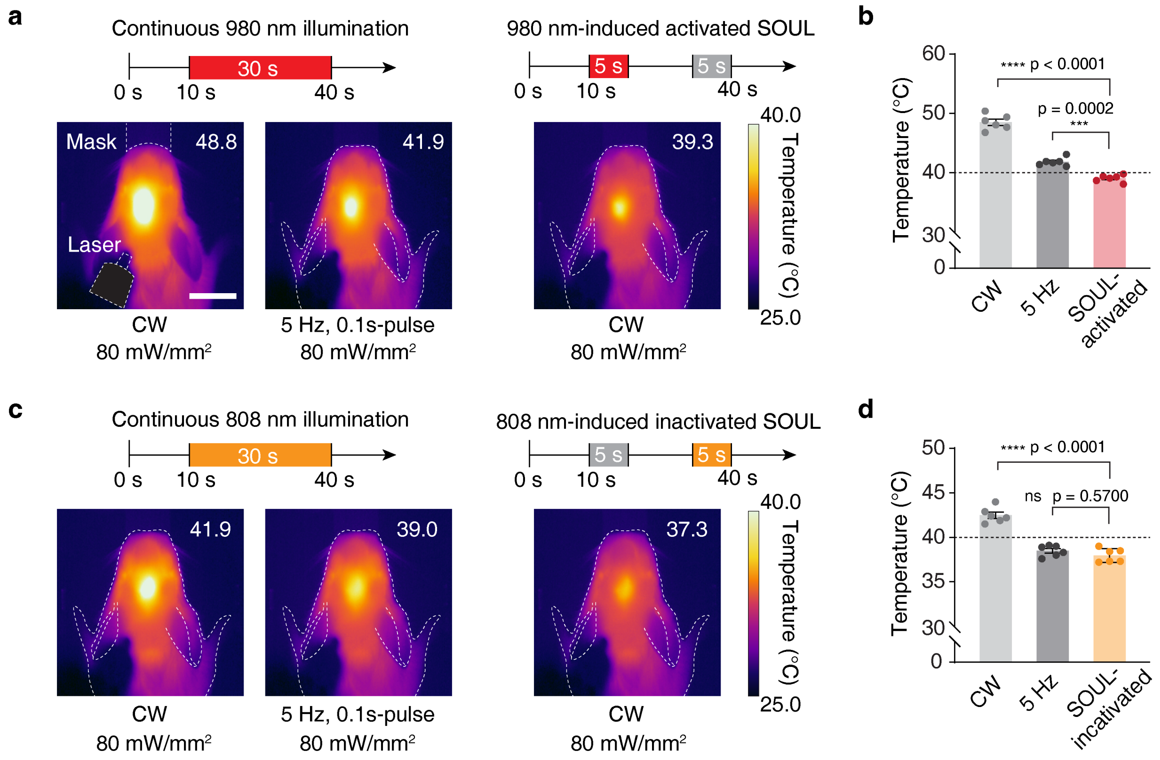

**Figure S20. Thermal effect comparison between different NIR illumination patterns at same wavelength.** (**a**) and (**c**) Thermal images showing the maximum temperature upon 980 nm and 808 nm illumination, respectively. The maximum temperature value is indicated in the upper right corner of the image. Illumination patterns and parameters are shown at the top and bottom of images. Scale bar, 1 cm. (**b**) and (**d**) Quantification of temperature for different illumination patterns. Data are shown as mean ± s.e.m. (n = 6 mice in each group, one-way ANOVA with Tukey's multiple comparison test). Dashed white lines indicate the mask, laser output and outline of mice.

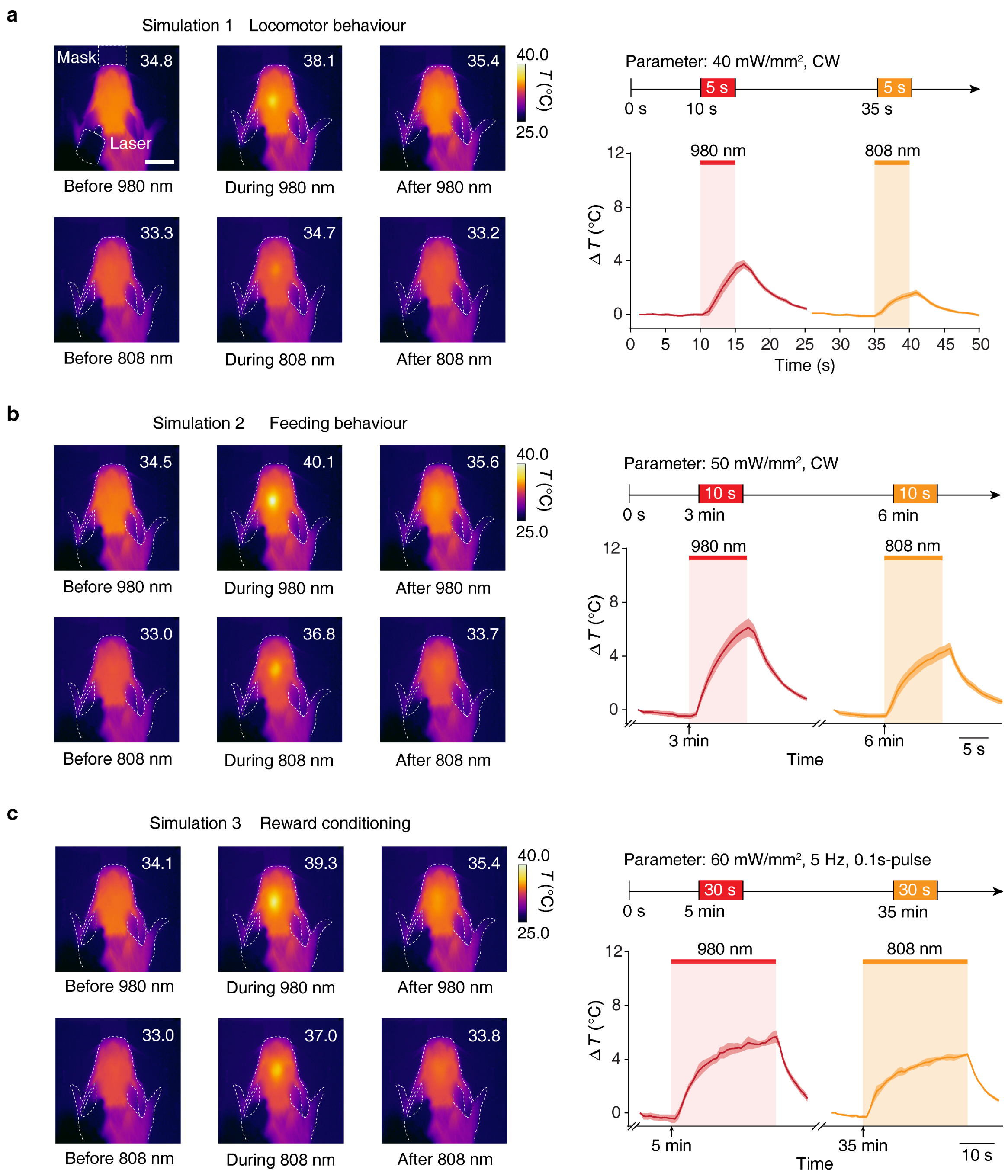

**Figure S21. Thermal effect simulations of different NIR illumination patterns applied in the behavioural paradigms.** Left, Representative thermal images acquired under different NIR illumination patterns and corresponding states. The maximum temperature value is indicated in the upper right corner of the image. All representative images are derived from the same mouse. Dashed white lines indicate the mask, laser output and outline of mice. Scale bar, 1 cm. Right, Temperature change dynamics of different NIR illumination patterns. The timeline and parameters of the simulated illumination pattern is displayed on the right. Red and orange shadows indicate the 980 nm and 808 nm illumination time window, respectively. Data are shown as mean ± s.e.m. (n = 6 mice in each group), the Time-axis scale is indicated at the bottom right of the respective data plot.

### Supporting Tables

**Table S1. Summary of complete statistical annotation.**

| Figure number | Detailed p-value |
|---|---|
| Figure 3 | For (+)/(-)/(-): Pre vs. During, p = 0.8475; vs. Post, p = 0.7737; During vs. Post, p = 0.4503.<br>For (+)/(+)/(-): Pre vs. During, p = 0.4201; vs. Post, p = 0.4362; During vs. Post, p = 0.8722.<br>For (+)/(-)/(+): Pre vs. During, p = 0.9727; vs. Post, p = 0.6973; During vs. Post, p = 0.6474.<br>For (+)/(+)/(+): Pre vs. Post, p = 0.8475; vs. Post, p = 0.8543. |
| Figure 4 | For (+)/(-)/(-): Pre vs. During, p = 0.6667; vs. Post, p = 0.6851; During vs. Post, p = 0.4242.<br>For (+)/(+)/(-): Pre vs. During, p = 0.9720; vs. Post, p = 0.9825; During vs. Post, p > 0.9999.<br>For (+)/(-)/(+): Pre vs. During, p = 0.1066; vs. Post, p = 0.43783; During vs. Post, p = 0.9654.<br>For (+)/(+)/(+): Pre vs. Post, p = 0.8475; vs. Post, p = 0.4930. |
| Figure 5 | Pre-test vs. Post-test: (+)/(-)/(-), p = 0.6606; (+)/(+)/(-), p = 0.1092; (+)/(-)/(+), p = 0. 4727. |
| Figure S13 | (+)/(+)/(+) vs. (+)/(-)/(-), p = 0.9551; vs. (+)/(+)/(-), p = 0.9891; vs. (+)/(-)/(+), p = 0.9638. |
| Figure S17a | Pre-test: (+)/(+)/(+) vs. (+)/(-)/(-), p = 0.9446; vs. (+)/(+)/(-), p = 0.2042; vs. (+)/(-)/(+), p = 0.9988.<br>Post-test: (+)/(+)/(+) vs. (+)/(-)/(-), p = 0.0995; vs. (+)/(+)/(-), p = 0.1253; vs. (+)/(-)/(+), p = 7225. |
| Figure S17b | Pre-test: (+)/(+)/(+) vs. (+)/(-)/(-), p = 0.9883; vs. (+)/(+)/(-), p = 0.1188; vs. (+)/(-)/(+), p = 0.9178.<br>Post-test: (+)/(+)/(+) vs. (+)/(-)/(-), p = 0.0937; vs. (+)/(+)/(-), p = 0.0937; vs. (+)/(-)/(+), p = 0.4582. |

#### Supporting Videos

**Video S1**. Video of a complete trial for NIR+/odUCNPs+/SOUL+ group mice in locomotion enhancement test related to the M2 brain region.

**Video S2**. Video of a complete trial for NIR+/odUCNPs+/SOUL+ group mice in free-accessing feeding test related to the LHA brain region.
